# A novel combinatorial treatment for Neurofibromatosis type 1 tumours revealed through cross-species genetic analysis

**DOI:** 10.1101/2025.05.06.652376

**Authors:** Megan Stevens, Seb Oltean, Benjamin E. Housden

## Abstract

Neurofibromatosis type 1 (NF1) is a multisystem genetic disorder associated with a wide range of symptoms, including the formation of tumours along nerves. These tumours can affect nerve function, leading to pain and loss of sensory or motor capacity. Drug treatments are currently limited, and relatively few new targets have emerged for NF1 tumors despite intensive research. Therefore, new drugs to target *NF1*-deficient tumour cells are urgently required. Here, we have used a cross-species screening strategy combined with bioinformatic analysis to identify candidates with a high chance of successful translation to humans. First, we used data from a synthetic lethal screen performed in *Drosophila* cells to identify novel drug targets of *NF1*. Then, we performed statistical enrichment analysis of the screen results to identify further candidate targets that were not conserved between *Drosophila* and humans and could not have been found in the screen. We identified and validated *hTERT* as a synthetic lethal partner gene to NF1 with high potential as a therapeutic drug target, using the FDA-approved inhibitor azidothymidine (AZT). The lethal effect of AZT was validated in peripheral neurofibroma *NF1*-deficient human cell lines and *NF1*-deficient malignant peripheral nerve sheath tumour cells. The effect of AZT was also conserved *in vivo,* highlighting the translational potential; treatment of mice with AZT inhibited *NF1* mutant xenograft growth to the same extent as the current clinically approved MEK inhibitor, selumetinib. Finally, we show that combined treatment with AZT and selumetinib resulted in a further synergistic reduction in *NF1*-deficient human cell viability. In conclusion, AZT represents a promising novel therapeutic for the treatment of *NF1* tumours, either alone or in combination with selumetinib.

## Introduction

Neurofibromatosis type 1 (NF1) is a relatively common genetic disease occurring in approximately 1 in every 3000 live births (1, 2). During childhood, it is associated with the formation of benign tumours, called neurofibromas, along peripheral nerves in addition to skeletal abnormalities, skin disfigurements and a range of other symptoms. Later in life, plexiform neurofibromas (PNs) can develop into malignant peripheral nerve sheath tumours (MPNSTs), which are frequently fatal (3).

NF1 is caused by the loss of one copy of the *NF1* gene, which can be either through inheritance or sporadic mutation. *NF1* is a tumour suppressor gene; the NF1 protein functions as a RAS GTPase Activating Protein (RASGAP) for H-, K-, N-RAS and R-RAS1, 2, and 3 (4-7). As a RASGAP, NF1 promotes the conversion of active RAS-GTP into its inactive form, RAS-GDP (8). Loss of one copy of *NF1* does not result in tumours; however, individual cells frequently lose their remaining copy of *NF1* through random mutation. Complete loss of *NF1* results in aberrant RAS signalling, and dysregulation of the pathways downstream of RAS, including the RAF/MEK/ERK and PI3K/AKT/mTOR pathways, which drives the formation of PNs. Furthermore, the acquisition of additional mutations can result in the progression to the potentially fatal MPNSTs (8).

It is not yet clear which of the numerous pathways downstream of RAS are required for the development of PNs. There has been much research targeting different components of the downstream RAS pathway, such as MEK and ERK (8); however, because RAS undergoes highly robust regulation, such therapies may not be ideal in the treatment of PNs. Due to RAS being implicated in many signalling pathways, chronic inhibition of RAS may not be an appropriate strategy for treating benign PNs. In addition, the picomolar affinity of RAS for GTP and the lack of suitable binding pockets for small molecules targeting RAS have resulted in RAS becoming an “undruggable” target (9). At present, only two drugs have been approved for the treatment of a subset of paediatric patients with PNs, the MEK inhibitors selumetinib and, very recently, mirdametinib. However, not all tumours are responsive to treatment, and serious side effects have been associated with the administration of MEK inhibitors (10-13). Various other targets have also been assessed, such as mTOR, receptor tyrosine kinases (RTKs), and RAF; however, the associated therapeutic strategies have often been found to be ineffective or too aggressive for long-term use (14). Therefore, new drugs that specifically target *NF1*-deficient cells, either alone or in combination with MEK inhibitors, are urgently needed.

In our previous study, we performed a genome-wide screen in *NF1-*deficient *Drosophila* cells to identify genes with a synthetic lethal interaction with *NF1 (15)*. Synthetic lethal partner genes can be exploited to exclusively kill, in this case, *NF1*-deficient cells. This is an attractive approach to cancer therapeutics because treatment is expected to be lethal to tumour cells but leaves wild-type, healthy cells viable (16). However, the use of synthetic lethality as a therapeutic strategy to treat tumours has so far resulted in few drugs successfully progressing to clinical trials (17). In fact, in the last decade, only PARP inhibitors, exploiting the synthetic lethal interaction between *PARP* and *BRCA1/2*, have succeeded in the clinic (18). One of the key factors associated with this lack of success is the lack of consistency between interactions identified in different genetic backgrounds, resulting in a lack of translation between model systems (19). Therefore, we used *Drosophila* cells to initially identify synthetic lethal interactions with genes mutated in *NF1* tumours. The conserved candidate interactions could be assessed in various model systems, including human cells, removing interactions specific to a single model system. This approach has previously proven to be successful in identifying mizoribine and palbociclib as promising candidates for the treatment of tuberous sclerosis complex (TSC) and Von Hippel-Lindau (VHL)-linked cancers, respectively (20-22), and in identifying chloroquine as a potential candidate for the treatment of NF1 (15). In all cases, hits from *Drosophila* synthetic lethal screens were validated with a high success rate in both human cells and mouse models.

One limitation of RNAi screens performed in *Drosophila* cells is that it is not possible to directly identify drug targets that are not conserved between *Drosophila* and humans. A second limitation is the level of noise associated with high-throughput screens, which can lead to false-negative results. In this study, we have addressed these limitations by performing statistical enrichment analysis of the list of *Drosophila* genes identified to have a synthetic lethal interaction with *NF1* to discover non-conserved genes, as well as genes that were missed in the screen that are likely to have a synthetic lethal interaction with *NF1* in humans. Computational prediction models have been widely used to complement wet-lab-based methods in identifying synthetic lethal pairs. Such methods include statistical-based methods, network-based methods, classical machine learning methods, and deep learning methods, which have been based on human and cross-species screens, as summarised in (23).

We applied statistical enrichment analysis of the list of *Drosophila* genes identified to have a synthetic lethal interaction with *NF1.* This approach proved effective, and we identified two new candidate drug targets that are not conserved between humans and *Drosophila*. Further investigation ultimately revealed that inhibition of the synthetic lethal partner gene *hTERT* with azidothymidine (AZT; a telomerase inhibitor) resulted in the induction of *NF1*-deficient cell death, both in a panel of human *NF1*-deficient cell lines, as well as in a mouse xenograft model. The mechanism of action was indicated to be complex, being telomerase-independent and additionally involving the HSP90 proteins. Furthermore, combined treatment with AZT and selumetinib resulted in synergistic cell death of *NF1* tumour cells. This work demonstrates that genetic screens performed in *Drosophila* can be used to identify candidate therapies targeting proteins that are not conserved between *Drosophila* and humans. In addition, we have identified a novel drug combination with the potential to effectively treat NF1 tumours.

## Methods

### Statistical enrichment analysis

Statistical enrichment analysis was used to identify additional candidates to those identified in the initial screen performed in our previous study (15). Gene ontology (GO) and semantic similarity analysis were performed using the GOSemSim package in R (24) and the online Revigo tool (http://revigo.irb.hr/). KEGG pathway analysis was performed using the KEGG Pathway Database (https://www.genome.jp/kegg/pathway.html). In addition, STRING was used to perform protein– protein interaction functional enrichment analysis (https://string-db.org/).

### Cell culture

*Drosophila* Schneider (S2R+) cells, both WT and dNF1-KO, were cultured at 25°C in Schneider’s media (Gibco) containing 1% antibiotic (Gibco) and 10% foetal bovine serum (Gibco). The following human cell lines were used: C8 *NF1^+/-^* and C23 *NF1^-/-^* immortalized human Schwann cell (SC) lines, derived from the ipn02.3 2λ cell line using CRISPR/Cas9 gene editing (25); two immortalized human SCs derived from plexiform neurofibromas from NF1 patients, which included ipnNF95.11C (*NF^+/-^*) and ipNF95.11b ‘C’ (*NF1^-/-^*) cells (germline *NF1* mutation: c.1756delACTA), and ipNF09.4 (*NF1^+/-^*) and ipNF05.5 (*NF1^-/-^*) cells (germline *NF1* mutation: c.3456_3457insA) (a generous gift from Prof. M. Wallace); and two malignant peripheral nerve sheath tumour cell lines, sNF96.2 (*NF1^-/-^*) and ST8814 (*NF1^-/-^*) (obtained from ATCC). All human cell lines were cultured at 37°C in 5% CO_2_ in DMEM media (Merck) containing 1% antibiotic (Gibco) and 10% foetal bovine serum (Gibco). For ipnNF09.4 and ipNF05.5 cells, the media was supplemented with 50 ng/ml neuregulin-1 (NRG-1) (Sigma).

### Variable dose analysis (VDA)

VDA assays were performed as described previously (26, 27). Briefly, on the day of transfection, S2R+ and dNF1-KO cells were plated at 1 × 10^4^ cells/100 µl culture media per well of a 96-well plate. Cells were incubated at 25°C for 40 minutes to allow adhesion. Cells were then transfected with 40 ng actin-GFP and 160 ng shRNA expression plasmid using 0.6 µl FuGENE^®^ HD transfection reagent (Promega) in a total volume of 10 µl. shRNA targeting *white* and *thread* were used as negative and positive controls, respectively. Plates were sealed and incubated for 4 days at 25°C in a humidifying chamber.

Flow cytometry was used to identify GFP-positive cells (transfection efficiency). The area under an inverted cumulative GFP distribution curve was used to measure cell viability, normalized to the positive and negative controls. (26)

### CellTiter-Glo assays

CellTiter-Glo assays (Promega; G7570) were used to measure cell population viability. These assays are based on a luminescent readout of ATP levels as a proxy for metabolically active cells. All cells were plated in white 384-well plates at a seeding density of 5 × 10^3^ cells/25 µl serum-free media. Before treatment, S2R+ cells were left to adhere for 40 min at 25°C, and human cells were left to adhere for 4 h at 37°C in 5% CO_2_. Cells in each well were treated with 250 nl of each drug (avapritinib, selumetinib, suramin, tazemetostat, mitoxantrone, erdafitinib, VLX1570, L-thyroxine, atovaquone, AZT, enzalutamide, BIBR-1532) in PBS or DMSO at varying concentrations using the Mosquito LV Genomics (SPT Labtech, Melbourn, Cambridgeshire, United Kingdom) and incubated for 48 h. CellTiter-Glo assays were performed per the manufacturer’s instructions, and luminescence was measured using a plate reader (TECAN Infinite M200 Pro, Männedorf, Switzerland).

### Cell count assays

Human cells were seeded in 24-well plates at a density of 2.5 × 10^4^ cells per well in complete culture media. After leaving the cells to adhere for 4–6 h at 37°C in 5% CO_2_, they were treated with AZT or selumetinib in DMSO in serum-free media. Counts were performed 24 and 48 h later. Trypan blue staining was used to exclude dead cells.

### Caspase assay

A caspase activity assay kit (Abcam; ab112130) assessed generic caspase activity. Cells were plated in 96-well plates in serum-free media and left to adhere for 4 h at 37°C in 5% CO_2_. Cells were subsequently treated with 250 µM per well of AZT and incubated for 48 h. The caspase assay was then performed per the manufacturer’s instructions. Briefly, the cell-permeable and non-toxic TF2-VAD-FMK fluorescent indicator, which irreversibly binds to caspase-1, -3, -4, -5, -6, -7, -8, and -9 in apoptotic cells, was added to the cell media for 4 h at room temperature. Cells were subsequently washed three times in PBS. Fluorescence intensity was measured using a plate reader at Ex/Em 490/525 nm (TECAN Infinite M200 Pro). To account for changes in cell density, caspase fluorescence was normalised to DAPI fluorescence in each well (Ex/Em 370/485 nm). Measurements in *NF1*-deficient cells were normalised to the average of control cells.

### NF1 tumour xenografts and toxicity assays

All experiments were conducted in accordance with UK legislation and with local ethics committee approval (University of Exeter AWERB) under the project license P983E59F3. Mice were housed in individually vented cages (Techniplast), with 3–4 mice per cage. They were housed in a sterile room with a 12 h/12 h light/dark cycle, 21 °C, and 45–55% humidity. Mice were handled under a laminar airflow hood and underwent procedures in a sterile surgical room. Mice were 6–8 weeks of age at the beginning of each study. Two million ST8814 (RRID: CVCL_8916) cells mixed 1:1 with Matrigel (ThermoFisher Scientific, Waltham, MA USA) were injected subcutaneously into the right flank of male CD-1 nude mice (Athymic Nude Mice obtained from Charles River (Crl:NU(NCr)-Foxn1nu)). Tumours were measured with a caliper three times weekly, and the tumour volume was calculated using the following formula: [(length + width) / 2] × length × width. Once the tumour diameter showed an increase over two separate measurements, mice were intraperitoneally injected with either 0.1% DMSO or AZT (100 mg/kg in 0.1% DMSO) or received selumetinib (25 mg/kg in saline) via oral gavage three times per week (n = 6 mice per group). Mice were culled by cervical dislocation (Schedule 1) when the control tumour sizes reached the allowed endpoint in the project licence (12 mm in diameter). The tumours were immediately dissected, flash-frozen, and stored at –80°C for further analysis.

Toxicity studies were performed using C57BL/6 Inbred Mice (Jax® Mice Strain) obtained from Charles River Laboratories.

### Reverse transcriptase-quantitative polymerase chain reaction (RT-qPCR)

RNA was extracted from cells and tissues using the Monarch Total RNA Miniprep Kit (New England Biolabs). One-step RT-qPCR was performed using the GoTaq® RT-qPCR System (Promega). The delta–delta Ct method was used to evaluate the normalized gene expression. Experiments were repeated with at least three biological repeats.

### Quantitative telomeric repeat amplification protocol (qTRAP)

To assess the telomere activity, we used the qTRAP method described by Jiang et al. (28). ST8814 cells were treated with various concentrations of AZT for 48 h followed by an assessment of telomerase activity.

### siRNA knockdown

DsiRNAs are chemically synthesized, 27 nt RNA duplexes available from IDT that are optimized for Dicer processing. ST8814 cells were transfected with a DsiRNA (20 nM) targeting hTERT (si- hTERT; hs.Ri.TERT.13.1: 5’-GGAAUCAGACAGCACUUGAAGAGGG-3’), HSP90AA1 (si- HSP90AA1.13.1: %’-GCAUGGAAGAAGUAGACUAAUCUCT-3’), HSP90AB1 (si- HSP90AB1.13.1: 5’-GGACAGUGGUAAAGAGCUGAAAATT-3’) or a scrambled siRNA control (sc-siRNA; all purchased from IDT) for 48–96 h using Lipofectamine RNAiMAX transfection reagent (ThermoFisher Scientific), per the manufacturer’s instructions. The level of knockdown was assessed using RT-qPCR.

### Western blotting (WB)

Protein lysates from tumour xenografts were prepared in RIPA buffer supplemented with a protease and phosphatase inhibitor cocktail (ThermoFisher Scientific). Denatured protein samples were run on mini-PROTEAN® TGX Stain Free™ pre-cast gels (4–15%, BIORAD), which allow for visualisation and accurate analysis of the total protein loaded for each sample using a Gel-Doc™ EZ (BIO-RAD) imaging system. Once the proteins had been transferred on to a PVDF membrane, total protein could be quantified. Membranes were blocked in blocking solution (BIORAD) before being probed with anti-PARP (9542; Cell Signaling Technology) diluted 1:1000 in blocking solution at 4°C overnight. After washing membranes in TBS-Tween (0.1%), the membranes were incubated in fluorescent secondary antibody (Li-cor goat anti-rabbit) diluted 1:10,000 in blocking solution for 2 h at room temperature before washing. Full-length and cleaved PARP were normalised to the loaded protein for each sample, as quantified by the Gel-Doc™ EZ imaging system (BIO-RAD). Experiments were repeated on at least three biological repeats, with the relevant controls run on the same blot.

### Drug synergy

To test whether the combined effects of AZT and selumetnib were additive or synergistic, the coefficient of drug interaction (CDI) was calculated for each cell line. The CDI was calculated as follows: CDI = AB/(A × B) (29-31), where A is the fold change of AZT compared to no-drug control, B is the fold change of selumetinib compared to no-drug control and AB is the fold change of the combined treatment compared to no-drug control. A CDI < 1 indicates synergy, CDI = 1 indicates additivity, and CDI > 1 indicates antagonism.

### Statistical analyses

Comparisons of two datasets were performed using either the Student’s t-test or the Mann– Whitney U test, depending on whether the data was normally distributed. When comparing three or more groups, a one-way analysis of variance (ANOVA) or Kruskal-Wallis test was performed, followed by post-hoc tests to assess multiple comparisons. In statistical analysis with two independent variables (i.e., time and drug), a two-way ANOVA was used with post-hoc testing for multiple comparisons. All statistical analyses were performed with GraphPad Prism, and a P-value < 0.05 was considered to be statistically significant.

## Results

### Using data from Drosophila genetic screens to identify non-conserved candidate drug targets relevant to humans

We previously reported the results of a genetic screen performed in *Drosophila* cells to identify genes that cause specific death of *NF1* mutant cells (15). This resulted in the discovery of 46 candidate drug targets (**Table S1**). While the use of genetic screens in *Drosophila* has proved to be an effective tool for drug target discovery, it is limited by the conservation of genes between *Drosophila* and humans. It is not possible to directly identify candidate drug targets that are not conserved between *Drosophila* and humans. In an attempt to overcome this issue, we searched for human genes with functional annotations that are enriched among the hits from our *Drosophila* genetic screen. We hypothesised that such genes are likely to have related functions to hits from the screens and, therefore, have potential as novel therapeutic targets for the treatment of *NF1*-deficient tumours. We also considered whether there were additional candidate targets in pathways identified by our initial screens, but they themselves were not identified as hits due to noise. We used a range of methods to perform statistical enrichment analysis of the 46 candidate genes, including GO-term enrichment and semantic similarity analysis, KEGG pathway enrichment, and further protein–protein interaction mapping (**Table S2–S9**). A flow diagram to show the process of enrichment analysis is shown in **Fig. 1A**.

**Figure 1.**
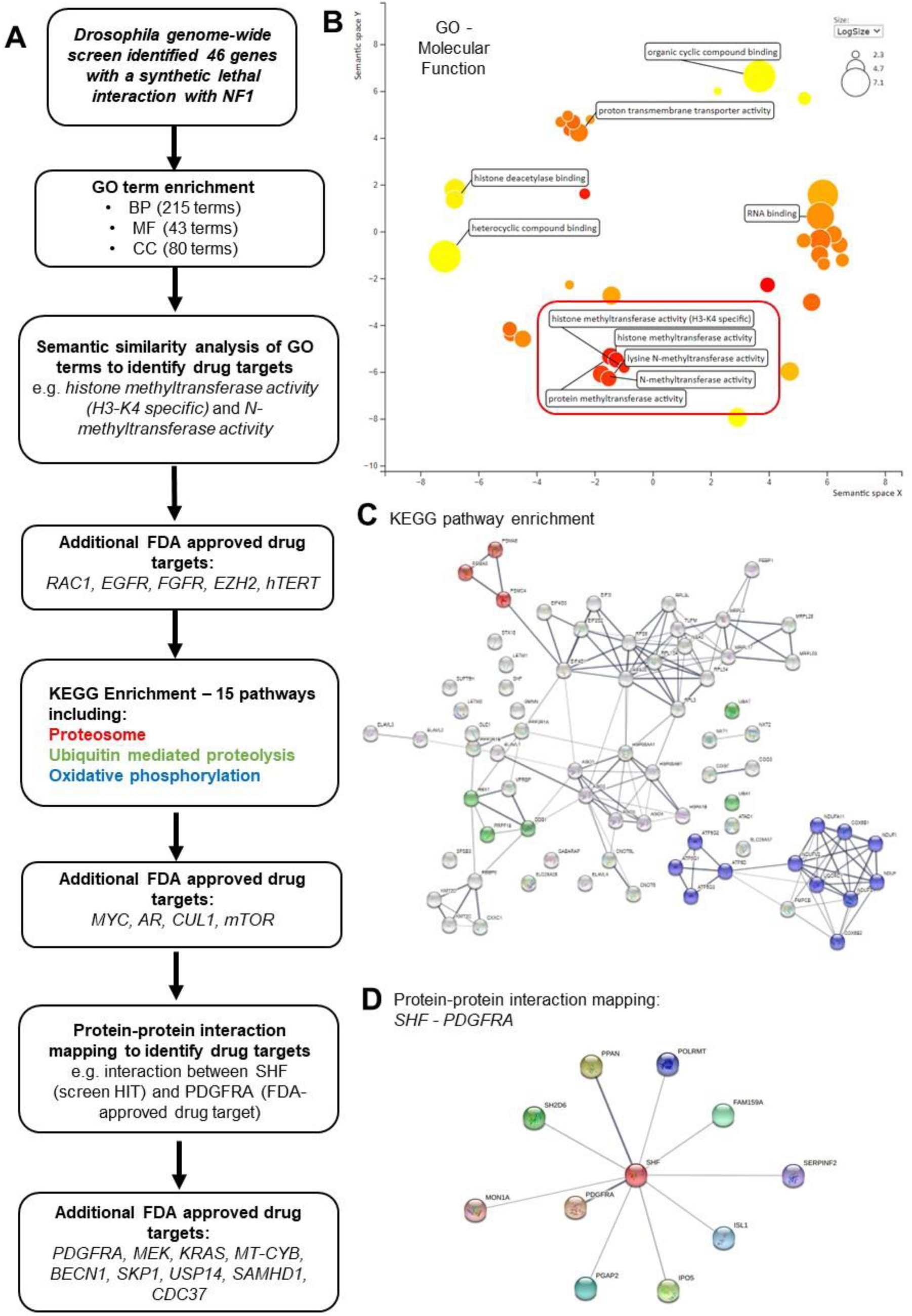
Statistical enrichment analysis of a *Drosophila* genome-wide screen to identify genes with synthetic lethal interactions with *NF1* that are druggable targets. **(A)** Flow diagram to show the process of enrichment analysis. As reported in our previous study, the *Drosophila* genome-wide screen identified 46 genes that have a synthetic lethal interaction with *NF1 (15)*. Throughout the enrichment analysis, we only selected genes that could serve as druggable targets. During the first stage, we identified GO terms that were significantly represented by the 46 candidate gene terms (FDR < 0.05). This resulted in 215 enriched terms in biological processes (BP), 43 enriched terms in molecular function (MF), and 79 enriched terms in cellular component (CC) subcategories. Semantic similarity between terms was visualised using the Revigo tool (http://revigo.irb.hr/), with an example shown in **(B)** for MF; semantic similarity was observed between “*histone methyltransferase activity (H3-K4 specific)”* and *“N-methyltransferase activity”*. Genes included in terms identified to have semantic similarity were screened for those that could be targeted by FDA-approved drugs, resulting in the addition of *RAC1, EGFR, FGFR, EZH2,* and *hTERT*. KEGG pathway enrichment analysis identified 15 pathways associated with the candidate gene list (three examples shown in **(C)**), which were used to screen for additional genes located upstream/downstream of the candidate gene signalling pathways that could be targeted with FDA-approved drugs. This resulted in the identification of *MYC, AR, CUL1,* and *mTOR* (in addition to *EGFR,* which was identified in the GO analysis). Finally, we assessed protein–protein interactions of the individual genes in the screen using the String database (https://string-db.org/) to identify interactions with additional genes that could be targeted with FDA-approved drugs, with one example shown in **(D)**. This resulted in the identification of *PDGFRA, MEK, KRAS, MT-CYB, BECN1, STIP1, SKP1, USP14, SAMHD1,* and *CDC37*.

### GO enrichment and semantic similarity analysis

During the first stage of analysis, we identified GO terms that were significantly represented by the 46 candidate gene terms (FDR < 0.05). This resulted in 215 enriched terms in biological processes (BP), 43 enriched terms in molecular function (MF), and 79 enriched terms in cellular component (CC) subcategories (**Table S2–S4**). Semantic similarity between terms was visualised using the Revigo tool (http://revigo.irb.hr/) (**Table S5–S7**), with an example shown in **Fig. 1B** for MF; semantic similarity was observed between “*histone methyltransferase activity (H3-K4 specific)”* and *“N-methyltransferase activity”*. The GOSemSim package in R (24) and Revigo tool were used to assign a semantic contribution factor between 0 and 1, where >0.8 indicates “in a relationship” and 0.6–0.8 indicates “part of a relationship”. Genes included in terms identified to have semantic similarity were screened for those that could be targeted by FDA-approved drugs, resulting in the identification of *RAC1, EGFR, FGFR, EZH2*, and *hTERT*. Full data is provided in the Supplementary Tables.

### KEGG pathway analysis

KEGG pathway enrichment analysis identified 15 pathways associated with the candidate gene list (**Table S8**; including the three examples shown in **Fig. 1C**: proteosome, ubiquitin-mediated proteolysis, and oxidative phosphorylation). In addition, there were two pathways that were common to the candidate genes and *NF1*: MAPK signalling pathway and EGFR tyrosine kinase inhibitor resistance pathway. These pathways were used to screen for additional genes located upstream/downstream of the candidate gene signalling pathways that could be targeted with FDA-approved drugs. This resulted in the identification of *MYC, AR, CUL1*, and *mTOR*.

### Protein–protein interaction analysis

Finally, we assessed protein–protein interactions of the individual genes in the screen using the String database (https://string-db.org/) to identify interactions with additional genes that could be targeted with FDA-approved drugs (**Table S9**). Some examples include the interaction of *SHF* (candidate gene) with *PDGFRA* (FDA-approved drug target) (**Fig. 1D**) and the interaction of *PEBP1* (candidate gene) with *MEK* and *KRAS* (both FDA-approved drug targets). This resulted in the identification of *PDGFRA, MEK, KRAS, MT-CYB, BECN1, SKP1, USP14, SAMHD1,* and *CDC37*.

In total, from the bioinformatic analysis, we identified 18 candidate human proteins that could be targeted with FDA-approved drugs, 16 of which had *Drosophila* orthologs and two of which did not.

### Validation of candidate synthetic lethal interactions using VDA

To assess the validity of the enrichment analysis, we tested whether the 16 candidate *Drosophila* genes had a synthetic lethal interaction with *NF1* using low throughput VDA assays. Two shRNA-expressing plasmids were generated targeting each of the 16 genes and were transfected into WT S2R+ and dNF1-KO cells. The shRNA showing the greatest viability effect in dNF1-KO cells for each gene is shown in **Fig. 2A**. Of the 16 genes, 14 showed a >10% reduction in viability in dNF1-KO cells compared to WT controls with a significant difference (**Fig. 2A**; P < 0.01). These results indicate the success of the enrichment analysis in identifying genes with a synthetic lethal interaction with *NF1*.

**Figure 2.**
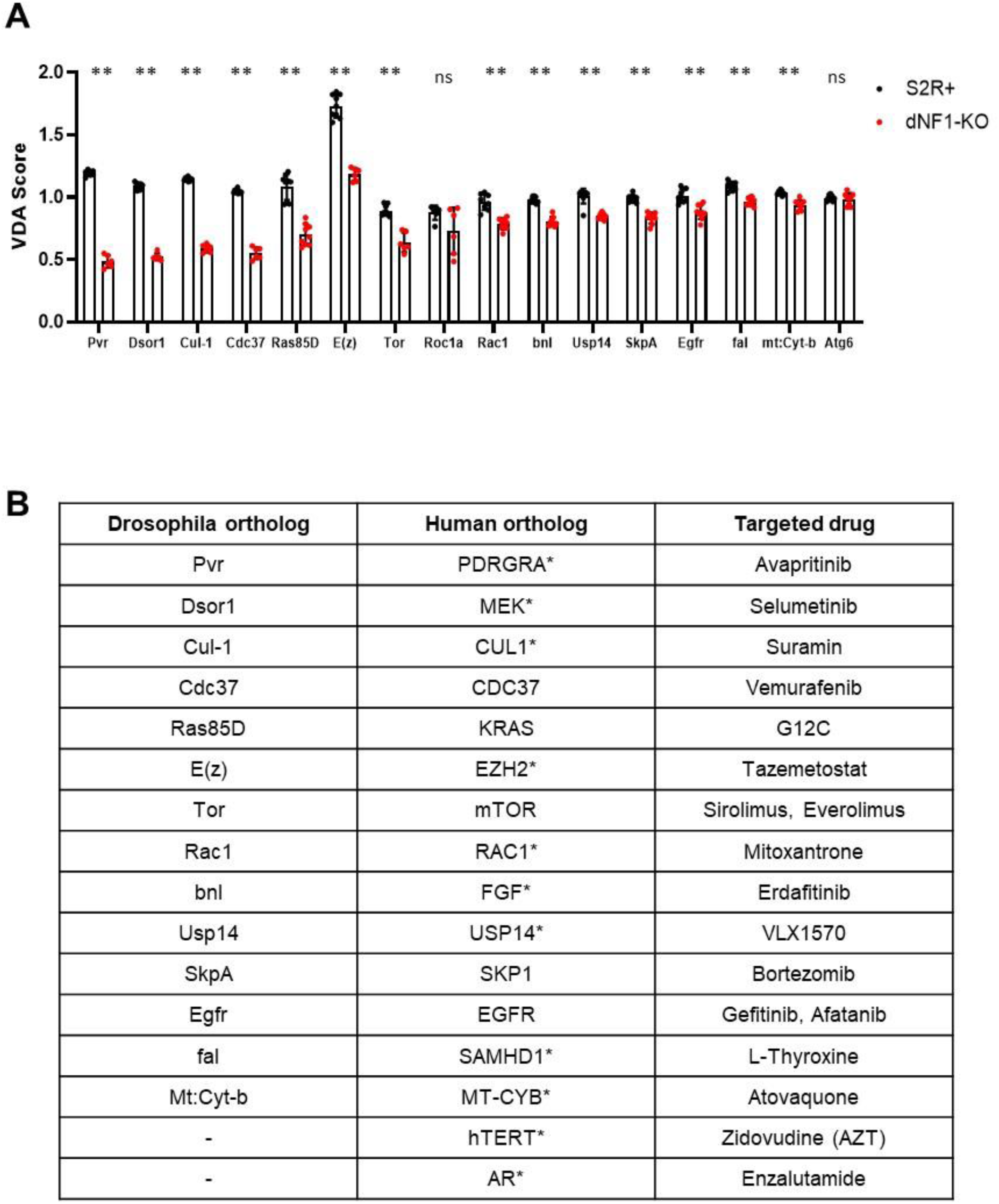
Validation of candidate genes from the statistical enrichment analysis. **(A)** VDA assays were performed for 16 candidate genes (with the exclusion of hTERT and AR as they do not have *Drosophila* orthologs), with two shRNAs per gene, in WT S2R+ and dNF1-KO cells. The shRNA with the greatest selective effect is shown in each case. shRNA knockdowns of candidate genes that resulted in a >10% reduction in dNF1-KO viability compared to S2R+ controls and were druggable targets are shown in red (n = 6–9, error bars indicate standard deviation, **P < 0.05 relative to S2R+ controls assessed using two-tailed, unpaired t-tests). **(B)** *Drosophila* hit genes, human orthologs, and inhibitors that can be used to target the synthetic lethal partner genes. * indicates genes/proteins that had not been previously assessed in the treatment of NF1; therefore, 11 drugs (which included selumetinib as a positive control) were taken forward to the next stage of the study.

Some of the genes that we found to have a synthetic lethal interaction with *NF1* have already been considered potential therapies to treat NF1 tumours. One example is *MEK*, which can be targeted with the FDA-approved drug selumetinib; however, not all tumours are responsive to treatment, and serious side effects have been associated with the administration of selumetinib (10-13). Another example is *EGFR*; however, reduced *NF1* expression has been found to result in resistance to EGFR tyrosine kinase inhibitors (32). *Dsor1* (*MEK*) and *Ras85D* (*KRAS*) were in the top five candidate drug targets (based on the difference in the viability effect in dNF1-KO and S2R+ cells using VDA assays), highlighting the validity of the assay. This indicates the high potential for this approach to identify quality targets for new therapies. To identify potential drugs for repurposing to treat NF1 tumours, we filtered the candidate gene list to those that have not been previously targeted in relation to NF1 therapy (**Fig. 2B**; * indicates novel target genes in addition to *MEK* as a positive control) and that could be inhibited with existing clinically approved drugs. The addition of the two human genes without *Drosophila* orthologs (*hTERT* and *AR*) resulted in 11 candidate genes.

### AZT selectively affects NF1-deficient human cells

To validate our results further and to identify existing drugs that may be repurposed to treat NF1 tumours, we tested the 11 drugs targeting the novel genes identified to have a synthetic lethal interaction with *NF1*. These were tested in a *Drosophila* cell model (WT and dNF1-KO S2R+ cells) and in a panel of human PN cell lines, including: C8 *NF1^+/-^* and C23 *NF1^-/-^* immortalized human SC lines, and two immortalized human SCs derived from PNs, ipnNF95.11C (*NF^+/-^*) and ipNF95.11b ‘C’ (*NF1^-/-^*) cells, and ipNF09.4 (*NF1^+/-^*) and ipNF05.5 (*NF1^-/-^*) cells (48 h treatment over a range of concentrations; cell viability assessed with the CellTiter Glo assay; **Fig. S1**). The drug found to have the most significant and consistent selective effect on *NF1*-deficient cell viability across the panel of cell lines was AZT (telomerase inhibitor) (**Fig 3A-B, Fig S1**).

**Figure 3.**
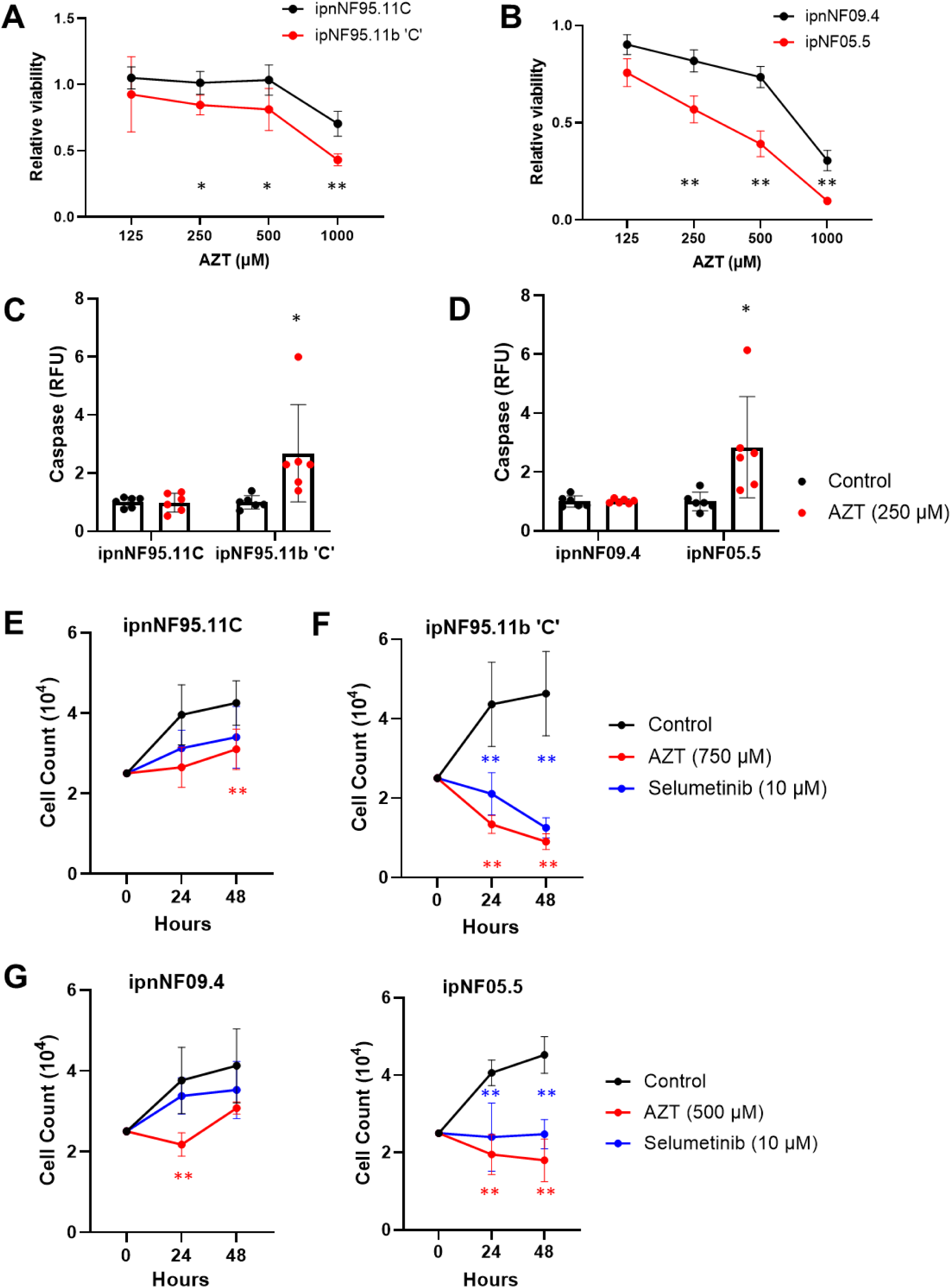
AZT selectively affects human *NF1*-deficient PN cells. After 48 h treatment in serum-free media, AZT significantly reduced *NF1^-/-^*cell viability relative to *NF1^+/-^* controls at varying doses in the PN cell lines ipnNF95.11C/ipNF95.11b ‘C’ **(A)** and ipnNF09.4/ipNF05.5 **(B)**, as measured with the CellTiter Glo assay (n = 8; P values obtained using a two-way ANOVA with Tukey for multiple comparisons; *P < 0.05, **P < 0.01). Furthermore, AZT (250 µM; 48 h in serum-free media) upregulated caspase activity in *NF1^-/-^* cells but not in *NF1^+/-^* control cells **(C, D)** (n = 3–6; P values obtained using the Student’s t-test; *P < 0.05 vs Control). Cell counts over 48 h revealed that selumetinib and AZT significantly reduced *NF1^-/-^* cell viability relative to *NF1^+/-^* controls in ipnNF95.11C/ipNF95.11b ‘C’ cells (n = 8 for Control, n = 4 for AZT and selumetinib) **(E, F)** and ipnNF09.4/ipNF05.5 cells (n = 8 for Control, n = 4 for AZT and selumetinib) **(G, H)** (P values obtained using a two-way ANOVA with Tukey for multiple comparisons; *P < 0.05, **P < 0.01). In all cases, data represent the mean and error bars indicate standard deviation.

AZT is an antiretroviral medication used to prevent and treat HIV/AIDS by acting as an inhibitor of HIV’s reverse transcriptase (33), the enzyme that the virus uses to make a DNA copy of its RNA. In human cells, AZT primarily functions to inhibit the activity of the telomerase, which is a highly specialised reverse transcriptase that maintains telomere length (34). *hTERT* does not have a *Drosophila* ortholog and was identified as a potential candidate gene in the GO and semantic similarity analysis of the genome-wide screen candidate genes (*HSP90AA1, HSP90AB1*). In each of the three human *NF1* mutant cell lines, AZT resulted in a significant reduction in *NF1^-/-^* cell viability relative to *NF1^+/-^* controls after 48 h treatment in serum-free media (**Fig. 3A and B, Fig. S2A**). The results of the CellTiter-Glo assays were validated by assessing caspase activity as a measure of cell apoptosis in response to AZT. After 48 h treatment, AZT (at the lowest dose determined to have a selective effect in each cell line) selectively and significantly increased caspase activity in both human PN *NF1*-deficient cell lines, which was not observed in the *NF1^+/-^* controls (**Fig. 3C and D**). Furthermore, AZT and selumetinib (at doses shown to have a selective effect in each cell line) inhibited cell proliferation in human PN *NF1*-deficient cell lines after 24 and 48 h treatment under serum-free conditions to a greater extent than *NF1^+/-^* controls (**Fig. 3E–H**). The most pronounced effect was observed at 48 h, with AZT treatment reducing the cell count relative to baseline (**Fig. 3F and H**). Although these doses appear to be fairly high for in vitro studies (250–750 µM), they are in line with previous studies assessing the effects of AZT on telomerase function and survival in human cells (35, 36).

Furthermore, the CellTiter-Glo assay was used to assess the effect of AZT on MPNST cell viability in comparison to SC WT control cells, indicating that AZT reduced *NF1* MPNST cell viability in sNF96.2 and ST8814 cells (**Fig. S2B and C**).

These results demonstrate that *NF1*-deficient cells can be selectively targeted with AZT, with the effects conserved across a panel of human *NF1* mutant cell lines. Therefore, AZT was identified as a candidate drug for the potential treatment of *NF1*-associated tumours.

### AZT reduced NF1-deficient MPNST tumour xenograft growth in vivo

Drug effects in cultured cells are not always reflective of *in vivo* effects. Therefore, we tested AZT in comparison with selumetinib using a xenograft model of *NF1-*deficient MPNST tumours. Although some NF1 PN tumour cells have been shown to form xenograft tumours, they are very slow growing and require signalling from *NF1^−/−^* peripheral nerves within the tumour microenvironment (37). On the other hand, highly aggressive MPNST *NF1^−/−^* cells grow rapidly in Matrigel xenografts, resulting in a highly reproducible preclinical model that allows the continuous quantitation of tumour growth during the study period; therefore, we chose to use this model to assess the effects on AZT on tumour growth. In mice implanted with ST8814 *NF1^-/-^*xenografts, once tumour growth had begun, we initiated treatment with AZT (100 mg/kg in 0.1% DMSO intraperitoneally, three times per week) or selumetinib (25 mg/kg in saline via oral gavage, three times per week). Using the FDA guidelines for dose conversion in mice, the recommended intravenous AZT dose in humans of 6 mg/kg/day (NICE guidelines) equates to 73.8 mg/kg/day in mice (6.0 × 12.3, which takes into consideration the Km factor, accounting for the different body surface areas); therefore, the chosen mouse dose of 100 mg/kg three times per week equates to 42.9 mg/kg/day, which is approximately within the range of the recommended intravenous dose in humans. Furthermore, using the same dose conversion, the selumetinib dose of 25 mg/kg three times weekly (10.7 mg/kg/day) is within the range of tolerated doses in humans (1.35–2.7 mg/kg/day, according to the NICE guidelines). Both drugs resulted in a significant reduction in tumour xenograft growth over a period of 3 weeks in comparison to vehicle-treated controls (**Fig. 4A**). Neither AZT nor selumetinib treatment was found to have toxic effects in these mice, which was also assessed in a prior study on C57BL/6 mice (**Fig. S3**). The selective effects of AZT were similar in strength compared to selumetinib, suggesting that AZT may represent a promising alternative to current therapeutic options.

**Figure 4.**
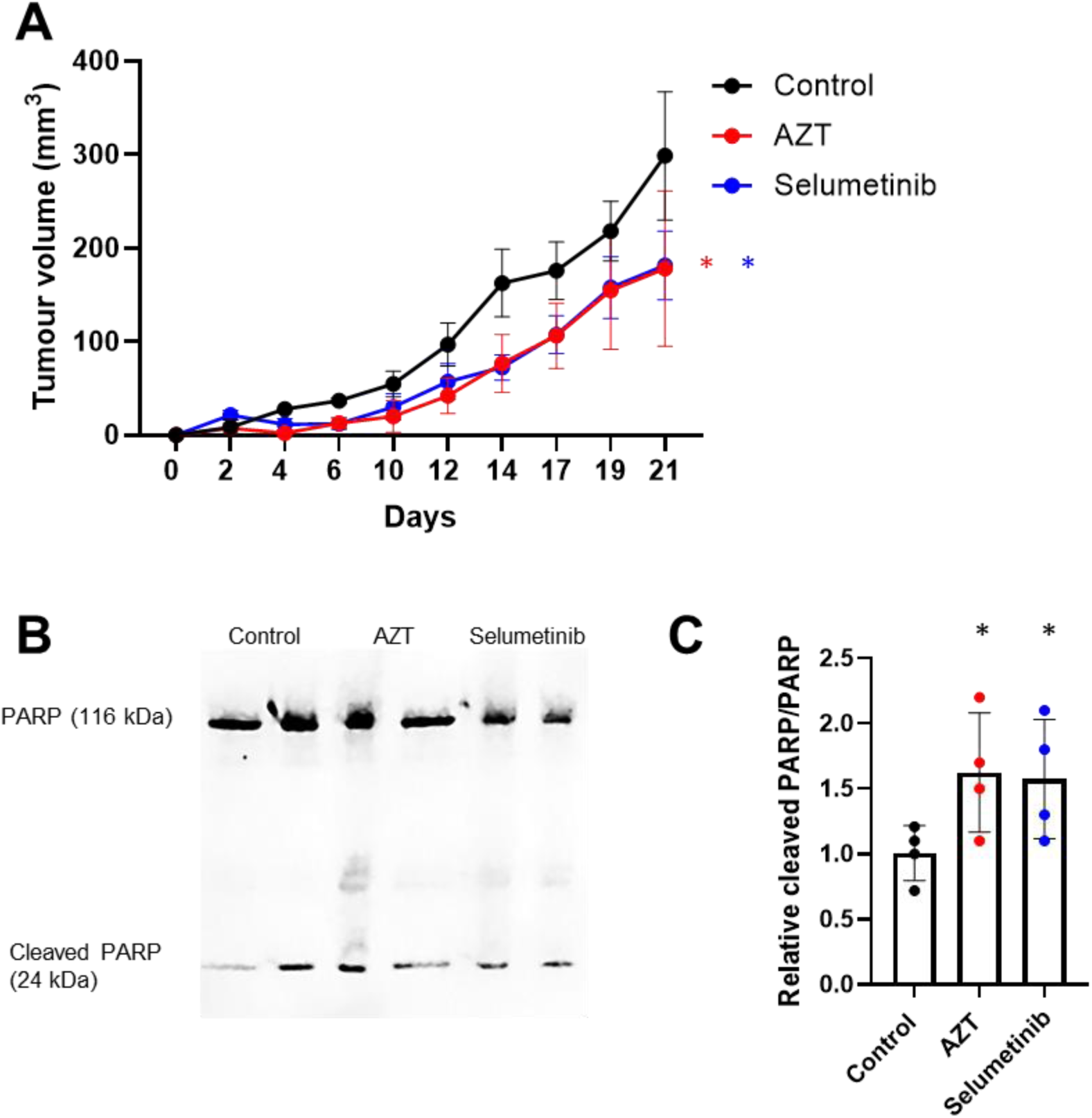
AZT inhibits *NF1-*deficient tumour xenograft growth *in vivo*. **(A)** In mice implanted with ST8814 *NF1^-/-^* xenografts, intraperitoneal injections of AZT (100 mg/kg, 3× weekly) or oral gavage with selumetinib (25 mg/kg, 3× weekly) significantly slowed tumour growth compared to vehicle-treated controls (n = 6; *P < 0.05 vs Control obtained using a two-way ANOVA). Error bars indicate the standard error of the mean. **(B)** Following extraction of the xenografts, western blotting revealed increased protein expression of cleaved PARP relative to PARP in AZT-treated mice in comparison to controls (n = 4; *P < 0.05 vs Control obtained using the unpaired Student’s t-test). Error bars indicate standard deviation.

AZT was found to induce *NF1*-deficient cell apoptosis *in vitro*; therefore, we used western blotting to assess the expression of a key apoptosis protein, cleaved PARP, in xenografts extracted from control, AZT, and selumetinib-treated mice. We found a significant increase in cleaved PARP/PARP in xenografts from AZT and selumetinib-treated mice, indicating that both resulted in cell apoptosis in the NF1 xenografts, thus slowing tumour growth (**Fig. 4B and C**).

### AZT may function via an hTERT-independent mechanism in NF1-deficient cells

As a structural analogue of thymidine, AZT primarily functions as an inhibitor of the reverse transcriptase enzyme. It is incorporated into the growing DNA chain by reverse transcriptase; however, it lacks a 3’ hydroxyl group necessary for further chain elongation, resulting in premature chain termination and halting viral replication (33). Although not usually a potent inhibitor of host cell DNA polymerases, at higher concentrations, AZT can be incorporated into the host DNA, resulting in damage to DNA synthesis (34). Therefore, we sought to determine the mechanism of action of AZT in *NF1-*deficient cells.

The telomerase complex consists of two main components, hTERC and hTERT. *hTERT* was the synthetic lethal partner gene identified in our enrichment analysis; however, upon initial testing of AZT in *NF1*-deficient Drosophila S2R+ cells, we saw no selective effect on viability (**Fig. S4**). This was hypothesised to be because there is no Drosophila ortholog for *hTERT*, leading us to suspect that the mechanism of action of AZT was through hTERT. Therefore, we assessed the effects of *hTERT* knockdown on the viability of ST8814 *NF1^-/-^* cells (a non-TERT immortalised cell line). A 71% knockdown in *hTERT* after 48 h resulted in a 22% reduction in cell viability (**Fig. 5A**). Furthermore, AZT resulted in a dose-dependent reduction in telomerase activity in ST8814 *NF1^-/-^* cells (**Fig. 5B**). However, we found no effect of AZT treatment (250 µM) for 48 h on *hTERT* mRNA expression in ipNF095.11b ‘C’ and ipNF05.5 *NF1^-/-^* cells (**Fig. 5C and D**). Furthermore, there were no significant changes in the mRNA expression of hTERT in the xenografts from mice treated with AZT compared to controls (**Fig. 5E**). To test whether the effect of AZT on *NF1^-/-^*- deficient cell viability was through hTERT, we treated the PN cells lines with a potent and selective hTERT inhibitor, BIBR-1532; we did not observe a selective effect on viability in either cell line (**Fig. 5F and G**). A similar finding was observed when we performed siRNA knockdown of *hTERT* in the ipnNF95.11C/ipNF95.11b ‘C’ and ipnNF09.4/ipNF05.5 paired cell lines. Although the mRNA expression of hTERT was higher in both *NF1^-/-^*-deficient cell lines (**Fig. Hi and Ii**), there was no selective effect on viability (**Fig. Hii and Iii**). Finally, ST8814 *NF1^-/-^* + *hTERT* knockdown cells were treated with varying doses of AZT (48 h knockdown + 48 h treatment with AZT). The effect of AZT on viability did not differ from the sc. siRNA control cells, indicating that the mechanism of AZT was independent of *hTERT* inhibition (**Fig. 5J**).

**Figure 5.**
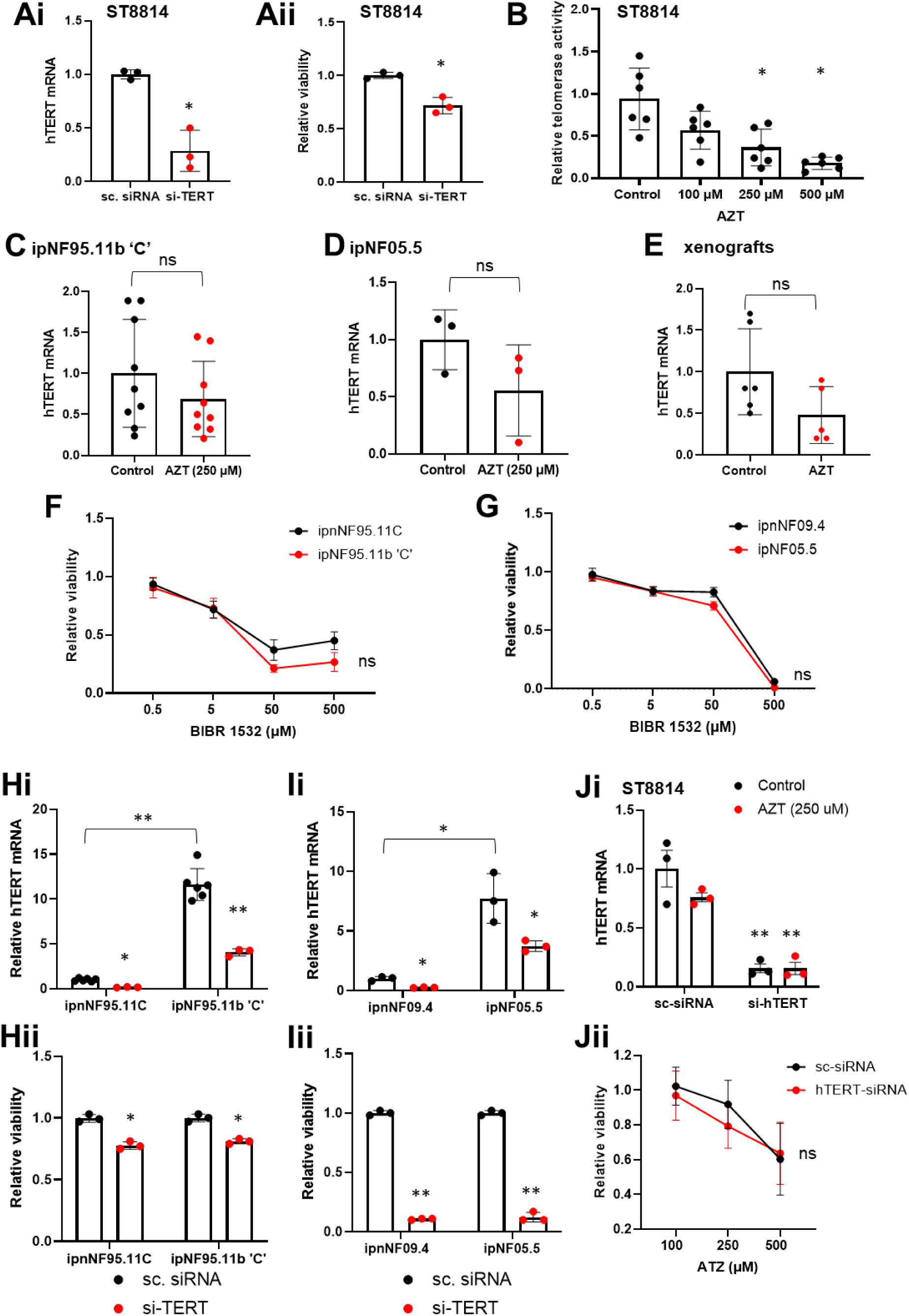
AZT may function through telomerase inhibition to exert its selective viability effect in *NF1-*deficient cells. **(A, B)** We performed siRNA-mediated knockdown of *hTERT* for 48 h in ST8814 *NF1^-/-^* cells **(Ai)**, which resulted in a reduction in cell viability **(Aii)**, as measured with the CellTiter Glo (n = 3; P values obtained using the Student’s t-test; *P < 0.05 vs sc. siRNA). **(B)** The qTRAP assay showed that AZT reduced telomerase activity in a dose-dependent manner (n = 6; P values obtained using the one-way ANOVA with Tukey post-hoc test; *P < 0.05 vs Control). However, AZT (250 µM) did not alter *hTERT* mRNA expression in ipNF95.11b ‘C’ *NF1^-/-^* cells **(C**; n = 9), ipNF05.5 *NF1^-/-^* cells **(D**; n = 3), or ST8814 *NF1^-/-^* xenografts **(E**; n = 5–6) (P values obtained using the Student’s t-test; P = ns vs Control). We used the CellTiter Glo assay to determine the effect of BIBR-1532, a potent and selective inhibitor of hTERT, in *NF1^-/-^* cells. After 48 h treatment with BIBR-1532 in SF media over a range of doses, we saw no selective effects on viability in the ipnNF95.11C/ipNF95.11b ‘C’ and ipnNF09.4/ipNF05.5 cells (n = 3–4; P = ns as assessed using a two-way ANOVA) **(F, G)**. The mRNA expression levels of *hTERT* were found to be significantly upregulated in ipNF95.11b ‘C’ and ipNF05.5 *NF1^-/-^* cells relative to *NF1^+/-^* controls (**Hi** and **Ii**; n = 3–6). Following siRNA mediated knockdown of *hTERT*, we saw a reduction in viability in both *NF1^+/-^* and *NF1^-/-^* cells, with no selective viability effect, as measured with the CellTiter Glo (**H** and **I**; n = 3; P values obtained using the Student’s t-test; *P < 0.05 and **P < 0.01 vs sc. siRNA). To determine whether AZT exerted its selective effect on *NF1^-/-^* cell viability independent of hTERT, we used an siRNA to knockdown *hTERT* in ST8814 *NF1^-/-^* MPNST cells for 96 h (n = 3; **P < 0.01 vs scrambled siRNA control, assessed using a Mann–Whitney test) **(F)**. Using the CellTiter Glo assay, we show that the selective viability effect of AZT (treated at 48–96 h) remained following the knockdown of *hTERT* (n = 6; ns = non-significant as assessed using a two-way ANOVA) **(G)**. Error bars indicate standard deviation.

Previous studies have shown that higher doses of AZT can affect DNA polymerases, causing mitochondrial dysfunction and cell cycle arrest (34, 36, 38-40). However, upon assessing the mRNA expression of several cell cycle-related genes following treatment of NF1-deficient cells with AZT for 48 h, we found that although some *NF1*-deficient cells have reduced expression of the cell cycle regulator p21, which was increased by AZT, the effects were not consistent across the panel of cell lines and genes tested (**Fig. S5**). In summary, these findings suggest that the selective effect of AZT is not functioning through DNA polymerase in *NF1*-deficient cells.

### AZT requires HSP90AA1 to alter the viability of some NF1-deficient cell lines

Going back to the enrichment analysis of the screen results, we found that *hTERT* clustered with *HSP90AA1* and *HSP90AB1* (both human orthologs of the screen hit *Hsp83*). Furthermore, these proteins play an important supporting role in the regulation and assembly of the telomerase complex (41, 42).

Both *HSP90AA1* and *HSP90AB1* expression were upregulated in the ipNF95.11b ‘C’ *NF1^-/-^* cells relative to ipnNF95.11C *NF1^+/-^* controls (**Fig. 6A**). However, this was not consistent across the PN cell lines. In ipNF05.5 *NF1^-/-^*cells, the expression of *HSP90AA1* was significantly lower, and *HSP90AB1* was the same compared to ipnNF09.4 *NF1^+/-^* controls (**Fig. 6B**). Furthermore, in the ipnNF95.11C/ipNF95.11b ‘C’ cells, siRNA-mediated knockdown of *HS90AA1* and *HSP90AB1* alone for 48 h did not alter the viability of *NF1^-/-^*cells, although a viability effect was observed in *NF1^+/-^* controls. However, dual knockdown of both *HSP90AA1* and *HSP90AB1* resulted in a viability reduction in both *NF1^-/-^* and *NF1^+/-^* cells (**Fig. 6Aiii**). In contrast, although knockdown of *HS90AA1* and *HSP90AB1* alone and in combination for 48 h altered the viability of ipnNF0.4 *NF1^+/-^* control cells, there was a further selective reduction in viability in ipNF05.5 *NF1^-/-^* cells with *HSP90AA1* and *HSP90AA1 + HSP90AB1* knockdown (**Fig. 6Biii**).

**Figure 6.**
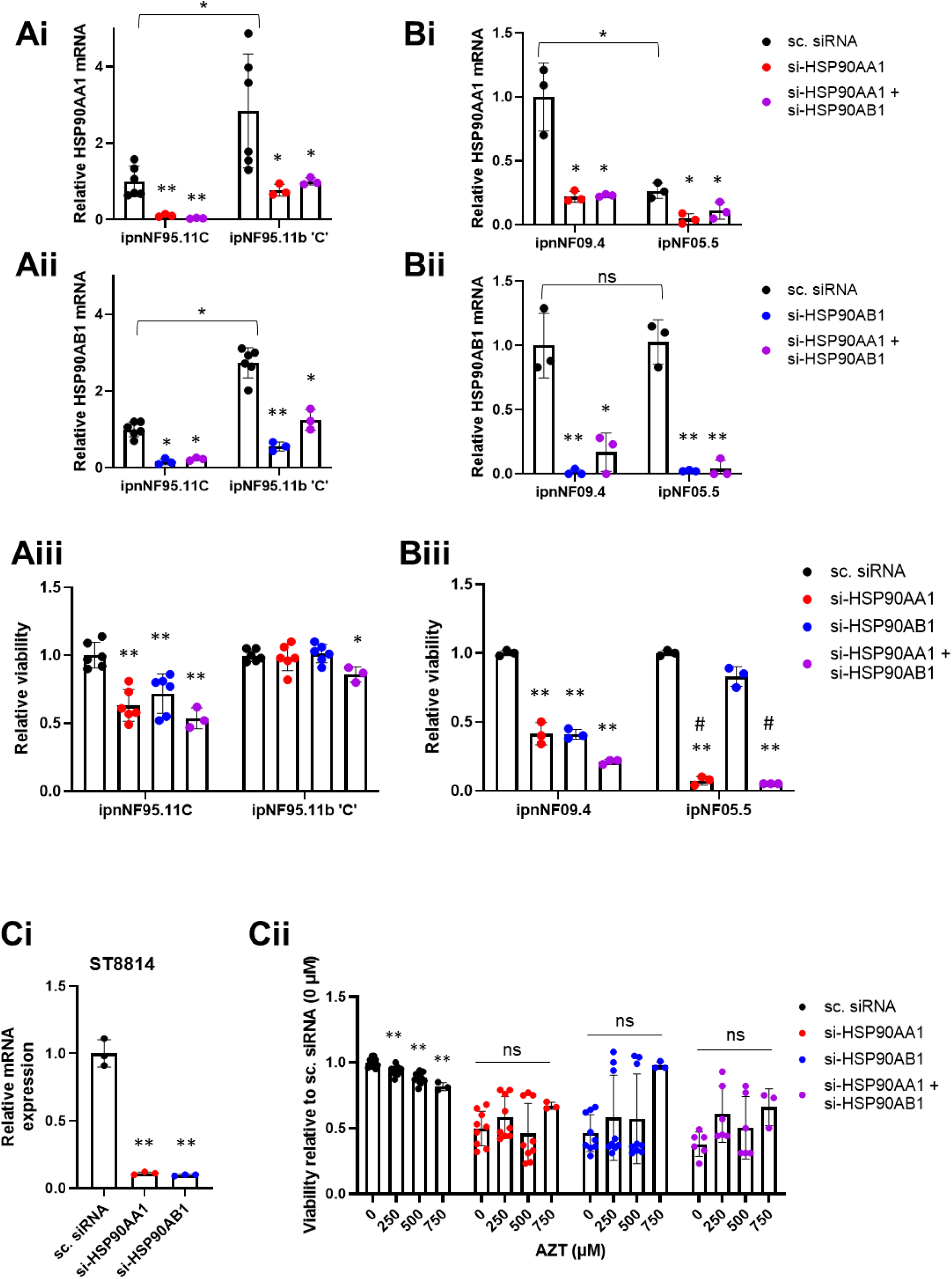
The effects of AZT on *NF1*-deficient cell viability are dependent on HSP90AA1. HSP90AA1 and HSP90AB1 are chaperone proteins that interact with and stabilise the telomerase complex. The mRNA expression levels of *HSP90AA1* and *HSP90AB1* were upregulated in ipNF95.11b ‘C’ *NF1^-/-^* cells relative to *NF1^+/-^* controls (**Ai** and **Aii**; n = 3–6; P values obtained using the Student’s t-test (for comparisons between cell lines) or the one-way ANOVA with Tukey post-hoc test (for comparisons within cell lines); *P < 0.05, **P < 0.01). However, siRNA-mediated knockdown of both genes for 48 h showed no effect on viability in ipNF95.11b ‘C’ *NF1^-/-^* cells, either alone or in combination, as measured with the CellTiter Glo assay (**Aiii**; P values obtained using one-way ANOVA with Tukey post-hoc test; **P < 0.01, *P < 0.05). In contrast, the mRNA expression level of *HSP90AA1* was downregulated in ipNF05.5 *NF1^-/-^* cells relative to *NF1^+/-^* controls **(Bi)**, with no change in *HSP90AB1* expression observed **(Bii;** n = 3; P values obtained using the Student’s t-test (for comparisons between cell lines) or the one-way ANOVA with Tukey post-hoc test (for comparisons within cell lines); *P < 0.05, **P < 0.01, ns = non-significant). Although *HSP90AA1* and *HSP90AB1* knockdown, alone and combined, resulted in a reduction in viability in ipnNF09.4 control cells, there was a further selective viability effect of *HSP90AA1* knockdown in ipNF05.5 *NF1^-/-^* cells, alone and in combination with *HSP90AB1* knockdown, as measured with the CellTiter Glo assay (**Biii**; n = 3; P values obtained using the Student’s t-test (for comparisons between cell lines) or the Kruskal-Wallis with post-hoc test (for comparisons within cell lines); **P < 0.01 sc. siRNA, ^#^P < 0.05 vs ipnNF0.94). (**C**) ST8814 cells underwent knockdown of *HSP90AA1/HSP90AB1* for 48 h followed by treatment with AZT for 48 h. Viability was assessed with the CellTiter Glo assay. Upon knockdown of *HSP90AA1* and *HSP90AB1* in ST8814 cells, we see that the dose-dependent effect of AZT on viability is no longer present. (n = 3 in Ci and n = 9 in Cii; P values obtained using one-way ANOVA with Tukey post-hoc test; **P < 0.01 vs sc. siRNA 0 µM AZT; ns = non-significant vs 0 µM AZT).

Finally, in NF1-deficient ST8814 cells with *HS90AA1*, *HSP90AB1,* or *HSP90AA1 + HSP90AB1* knockdown, the selective effects of AZT on cell viability were inhibited (**Fig. 6C**), indicating that AZT requires the presence of HSP90s to have an effect on *NF1*-deficient cell viability, at least in some *NF1*-deficient cell lines.

### Selumetinib enhances the viability effect of AZT

Selumetinib and mirdametinib, MEK1/2 inhibitors, are currently the only FDA-approved drugs for the treatment of tumours associated with neurofibromatosis. We treated human PN and *NF1^-/-^* MPNST cell lines with selumetinib (10 µM) +/- AZT (250 µM) for 48 h in serum-free media and performed CellTiter-Glo assays to measure cell viability (**Fig. 7**). When selumetinib was combined with AZT, we observed a further reduction in *NF1^-/-^,* but not *NF1^+/-^,* viability relative to AZT alone, indicating that selumetinib enhances the effects of AZT on *NF1^-/-^* viability.

**Figure 7.**
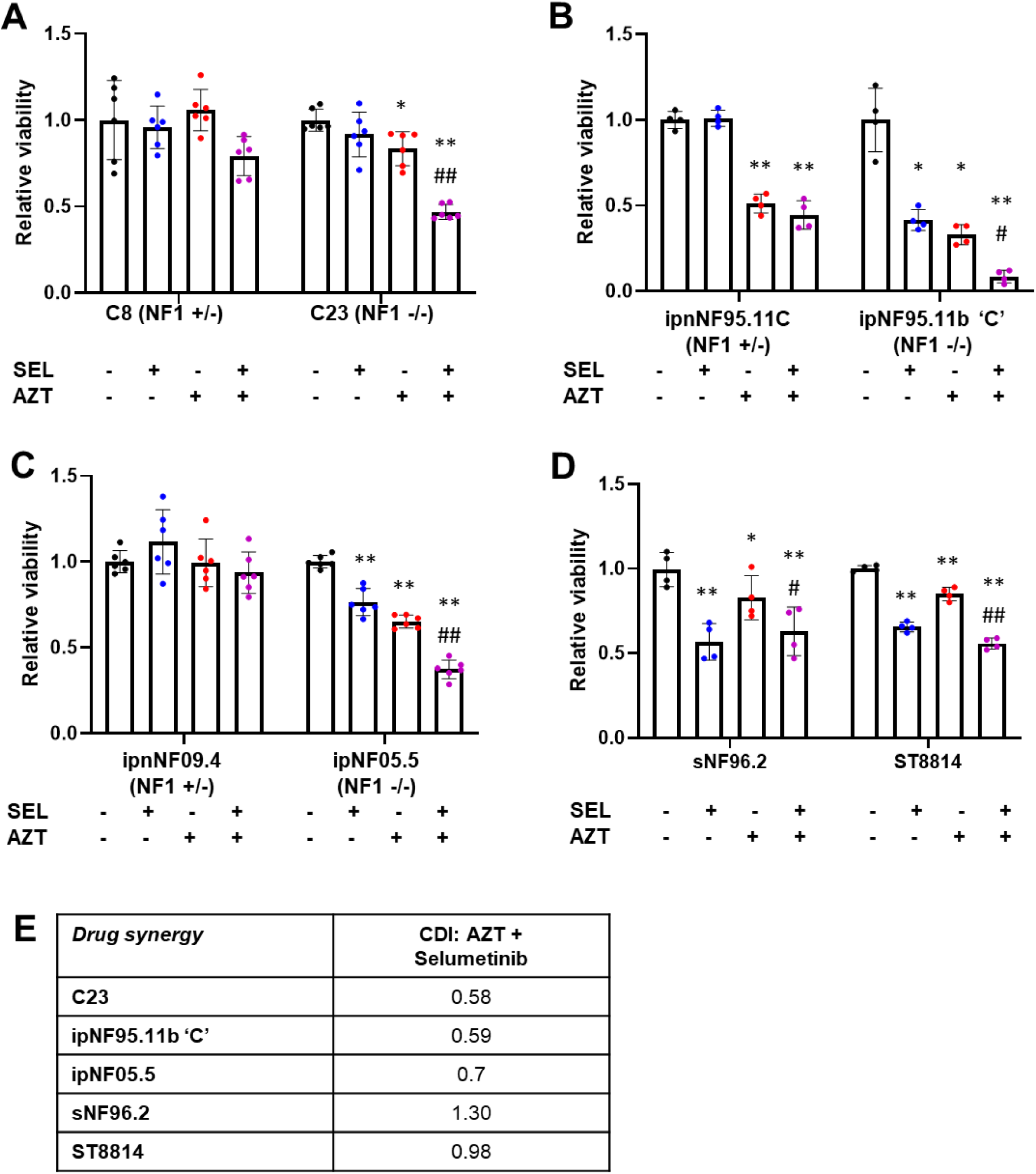
AZT and selumetinib synergise in PN-derived cells. **(A–C)** Human PN cells (C8/C23, ipnNF95.11C/ipNF95.11b ‘C’, and ipnNF09.4/ipNF05.5) and **(D)** *NF1^-/-^* MPNST cells (sNF96.2 and ST8814) were treated with 0 µM or 10 µM selumetinib combined with 0 µM or 250 µM AZT for 48 h. Alone, selumetinib significantly reduced the viability of ipNF95.11b ‘C’, ipNF05.5, sNF96.2, and ST8814 NF1^-/-^ cells, with no effect on the viability of C23 *NF1^-/-^* cells and *NF1^+/-^* controls, as measured with the CellTiter Glo assay. Similarly, AZT alone significantly reduced the viability of all *NF1^-/-^*cells, with a less extensive effect on ipnNF95.11C *NF1^+/-^* viability also observed. When used in combination, the selective viability effect in all *NF1^-/-^*cells was significantly enhanced (n = 4–6; P values obtained using a one-way ANOVA with Tukey for multiple comparisons; *P < 0.05 and **P < 0.01 vs untreated Control, ^#^P < 0.05 and ^##^P < 0.01 vs AZT alone). **(E)** Drug synergy between AZT and selumetinib was calculated using the coefficient of drug interaction (CDI). CDI < 1 indicates synergism, CDI = 1 indicates additivity, and CDI > 1 indicates antagonism.

To test whether these combined effects were additive or synergistic, the CDI was calculated for each cell line treated with AZT and selumetinib. We observed a relatively strong synergistic effect (CDI < 0.7) in all PN *NF1^-/-^* cells, with an additive/possible antagonistic effect observed in MPNST *NF1^-/-^* cells (**Fig. 6E**). These results suggest that AZT and selumetinib together may represent a powerful treatment for PN tumours associated with NF1.

## Discussion

This is a follow-up to our previous study that used dNF1-KO cells as a model to identify genes with synthetic lethal interactions with *NF1* using a genetic interaction screen (15). However, one limitation of RNAi screens performed in *Drosophila* cells is that it is not possible to directly identify drug targets that are not conserved between *Drosophila* and humans. A second limitation is that potential candidates could have been missed due to noise from the screen or ineffective reagents. Therefore, in the present study, to find such targets, we performed statistical enrichment analysis of the candidate genes to identify additional human genes that are likely to perform similar roles to the genes identified in the *Drosophila* screen. Furthermore, we used the enriched annotations to identify other proteins with functions related to the screen hits that might be more amenable to inhibition with existing FDA-approved drugs. As a result, we identified 18 more candidate genes (including two without *Drosophila* orthologs). We tested the 16 conserved genes using VDA and found that 14 had a synthetic lethal relationship with *NF1*. Drugs targeting these 14 proteins (plus two without *Drosophila* orthologs – 16 in total) were screened in a panel of human *NF1* mutant cell lines. The results showed that AZT (targeting telomerase) had the most consistent and selective viability effect in *NF1^-/-^* cells. Furthermore, *NF1* mutant xenografts from mice were decreased in size when mice were treated with AZT, indicating that AZT can slow *NF1* mutant MPNST xenograft growth *in vivo.* One of the suggested mechanisms of action of AZT in *NF1* mutant cells was telomerase inhibition associated with HSP90. However, the exact mechanism appears to be complex and may differ between *NF1*-deficient cell lines.

GO and KEGG pathway enrichment analysis of our list of genes identified two genes without fly orthologs to have a synthetic lethal interaction with *NF1*, *hTERT* and *AR*. Both have been targeted in anticancer therapies previously. AZT, a telomerase inhibitor, showed the most selective effect on viability in our panel of *NF1* mutant cell lines and was, therefore, taken forward to the next stage of the study.

The GO term “telomerase holoenzyme complex assembly” (GO:1905323) was significantly enriched in the GO enrichment analysis of the candidate gene list from the screen (FDR adjusted P value = 0.0059). AZT is an inhibitor of telomerase, which is a specialised reverse transcriptase (RT) that maintains telomeric length. *hTERT* encodes the functional catalytic component of the enzyme. The human holoenzyme telomerase is highly expressed in embryonic cells and is then repressed during adulthood; however, the enzyme is re-expressed in approximately 85% of solid tumours (43), which has prompted studies to inhibit telomerase with the HIV RT AZT. AZT binds to telomeres, inhibits telomerase, and promotes tumour cell apoptosis (35, 44). We found that in all three human PN *NF1^-/-^* cell lines and two *NF1^-/-^* MPNST cell lines, AZT resulted in a significant reduction in cell viability across a range of concentrations, with little effect on *NF1^+/-^* cells at concentrations up to 500 µM.

Following the success of AZT in selectively killing *NF1^-/-^*cells *in vitro,* we went on to assess the effect *in vivo*. We used the ST8814 *NF1^-/-^* MPNST xenograft mouse model because although some NF1 PN tumour cells have been shown to form xenograft tumours, they are very slow growing and require signalling from peripheral nerves within the tumour microenvironment (37).

When administered intraperitoneally three times per week over 3 weeks, AZT inhibited tumour growth at doses considered to be non-toxic over the study period, as has also been shown using the same dose in many previous xenograft studies (45). In addition, the dose we used (100 mg/kg) is reflective of the dose used *in vitro* (equating to 360 µM). The selective effects of AZT were similar in strength compared to selumetinib, suggesting that AZT may represent a promising alternative to current therapeutic options. The increased expression of cleaved PARP in xenografts from AZT-treated mice indicates that AZT is causing apoptosis of the *NF1*-deficient tumour cells, thus preventing tumour growth. AZT has been documented to induce apoptosis in many cancer cell lines (46, 47), as well as in tumour xenograft models(45). Its mechanism, inhibition of telomerase resulting in cell death, appears to be dependent on HSP90 in some *NF1^-/-^* cells, although the exact mechanism remains unclear.

The mechanism by which AZT selectively kills *NF1^-/-^*cells could be due to increased telomerase activity in *NF1*-deficient cells. Telomerase has been shown to be upregulated via portions of the RAS pathway (48). Indeed, we found that *hTERT* mRNA expression was higher in the two patient-derived PN *NF1^-/-^* cell lines in comparison to their paired *NF1^+/-^*controls. However, previous studies have shown that *hTERT* expression is not altered in *NF1-*deficient PNs or low grade MPNSTs (49). In addition, there have been reported examples in which AZT has selective effects on the viability of cancer cells that do not express hTERT or with no associated hTERT activity (46). Furthermore, we found no selective effects of AZT in Drosophila NF1-deficient cells, where telomere maintenance is not performed by the canonical telomerase and instead by a unique transposition mechanism (50). Therefore, it is likely that AZT acts, at least partially, via an uncharacterised mechanism to kill *NF1-*deficient cells. Using the qTRAP assay (28), we found that AZT was acting through a telomerase-dependent mechanism in the *NF1*-deficient MPSNT cell line ST8814. We used this cell line because the PN cell lines are TERT-immortalised and could, therefore, have altered sensitivity to the inhibition of telomerase activity. Therefore, although AZT showed inhibition of telomerase at these doses, it does not appear to be responsible for the selective effects on NF1 cells.

However, directly targeting hTERT, either through siRNA-mediated knockdown or treatment of cells with BIBR-1532, a potent and selective hTERT inhibitor, did not show a selective viability effect in any of the *NF1*-deficient cells. Furthermore, the viability effect of AZT remained in the presence of *hTERT* knockdown, and AZT had no effect on *hTERT* expression. These findings indicate that although AZT inhibits telomerase activity in *NF1*-deficient cells, this is independent of hTERT expression.

In the enrichment analysis of the screen results, *hTERT* was clustered with *HSP90AA1* and *HSP90AB1* (both human orthologs of the screen hit *Hsp83*). HSP90AA1 (Heat Shock Protein 90 Alpha Family Class A Member 1) and HSP90AB1 (Heat Shock Protein 90 Alpha Family Class B Member 1) are two isoforms of the molecular chaperone HSP90, a highly conserved protein that plays a crucial role in the proper folding, stability, and activation of a variety of client proteins, including those involved in signal transduction, cell cycle regulation, and stress responses. Furthermore, HSP90 is known to be associated with mTOR and AKT, and it plays an important role in telomerase activity (41, 42). HSP90AA1 and HSP90AB1 interact with the telomerase complex in the following ways: (1) Chaperoning of telomerase components, including TERT and TERC, and assisting in the assembly of the entire telomerase complex, which ultimately affects enzyme function. (2) Under certain stress conditions (such as heat shock or other cellular stresses), HSP90AA1 and HSP90AB1 help prevent the degradation of TERT, allowing telomerase activity to persist. Furthermore, in some cancers, for example, where telomerase is reactivated, HSP90 inhibitors have been investigated as potential therapeutic strategies to reduce telomerase activity and, thereby, limit cancer cell proliferation (51). For example, derivatives of Geldanamycin, a natural inhibitor of HSP90 isolated from *Streptomyces hygroscopicus*, have been shown to enhance the potency of telomerase inhibition by imetelstat in human osteosarcoma (52). Therefore, HSP90-mediated inhibition of telomerase could be a potential therapeutic option in NF1.

HSP90, AKT, TERT, S6K, and mTOR have been reported to form a physical complex (41), providing compelling evidence for mTOR-mediated control of telomerase activity. TERT requires interaction with both HSP90 and AKT for efficient telomerase activity (53, 54). Furthermore, HSP90 has been reported to stabilize the mTOR-TERT complex (55). It is suggested that this complex is necessary for cancer cell survival. In addition, the mTOR pathway has been reported to be tightly regulated by NF1, and mTOR inhibition with rapamycin resulted in a selective effect on NF1 (56). Interestingly, all of these components were hits in the synthetic lethality screen with NF1, highlighting the importance of telomerase activity in *NF1*-deficient cells with aberrant RAS signalling.

*HSP90AA1* and *HSP90AB1* encode Hsp90α and Hsp90β, respectively. These two isoforms are 85% identical; however, *HSP90AB1* is constitutively expressed and essential for development, whereas HSP90AA1 is stress-inducible (57). Both HSP90AA1 and HSP90AB1 are involved in stabilizing the telomerase complex. In this study, we found that the expression of *HSP90AA1* and *HSP90AB1* varied across different NF1 cell lines. In ipNF95.11b ‘C’ *NF1^-/-^* cells, the expression of both HSP90s was 2–3-fold higher than in ipnNF95.11C *NF1^+/-^* controls. As a result, the siRNA-mediated knockdown of *HSP90AA1* and *HSP90AB1* was less effective, possibly explaining why a selective effect on *NF1*-deficient cell viability was not observed. In contrast, the expression of *HSP90AA1* was 3-fold lower in ipNF05.5 *NF1^-/-^*cells compared to ipnNF09.4 *NF1^+/-^* controls. As a result, we saw a selective effect of *HSP90AA1* knockdown on ipNF05.5 *NF1^-/-^*cell viability. Furthermore, AZT did not result in a dose-dependent effect on ST8814 *NF1*-deficient viability in the presence of *HSP90AA1* and/or *HSP90AB1* knockdown. These findings show that AZT requires the presence of HSP90s to have an effect on *NF1*-deficient cell viability, at least in some *NF1*-deficient cell lines. Further studies are required to elucidate the roles of each HSP90 isoform in response to AZT treatment to selectively target *NF1*-deficient cells.

Finally, we tested the effects of combining AZT and selumetinib in NF1-mutant cell lines, as drug combinations can be more effective than either agent used alone. In addition, combinatorial treatments may allow the dose of each drug to be lowered, thus reducing the risk of any side effects. Particularly in the PN *NF1^-/-^*cell lines, we found that AZT strongly synergises with selumetinib (as indicated by a CDI < 0.7, which is deemed significant synergism), further highlighting the therapeutic potential of AZT in the treatment of NF1 tumours.

A future direction could be to test AZT in a NF1 PN-specific model, such as the *NF1^flox/flow^;PostnCre* genetically engineered model of PN (43), both alone and in combination with selumetinib. If successful, this could proceed to a clinical evaluation of AZT either alone or in combination with selumetinib as a potential new treatment for NF1 tumours.

## Conclusion

In conclusion, using multiple techniques, assays, and models, we have shown that targeting telomerase with the FDA-approved drug AZT has potential as a novel therapeutic strategy for the treatment of NF1 tumours. Further work is required to elucidate the mechanisms by which AZT is exerting this potential therapeutic effect.

## Acknowledgements

This study was funded by the MRC (grant no. MR/V009583/1) and an institutional CiC MRC award (parent award no. MR/X502820/1). We would also like to acknowledge Nerve Tumours UK for their support. For the purpose of open access, the author(s) has applied a Creative Commons attribution (CC BY) licence to any Author Accepted Manuscript version arising.

## Notes

### Competing Interest Statement

The authors have declared no competing interest.

