## Supplementary Figures for "A novel combinatorial treatment for Neurofibromatosis type 1 tumours revealed through cross-species genetic analysis"

**Fig. S1. Dose curves for each drug tested.** Using the CellTiter Glo assay, we performed dose curves to determine the effect of each drug on *Drosophila* (i) and human (ii) Nf1-deficient cell viability. (A) avapritinib; (B) selumetinib; (C) surmain; (D) tazemetostat; (E) mitoxantrone; (F) erdafitinib; (G) VLX1570; (H) L-thyroxine; (I) atovaquone; (J) enzalutamide; (K) AZT. Note that enzalutamide and AZT were only tested in human cells as there were no *Drosophila* orthologs.

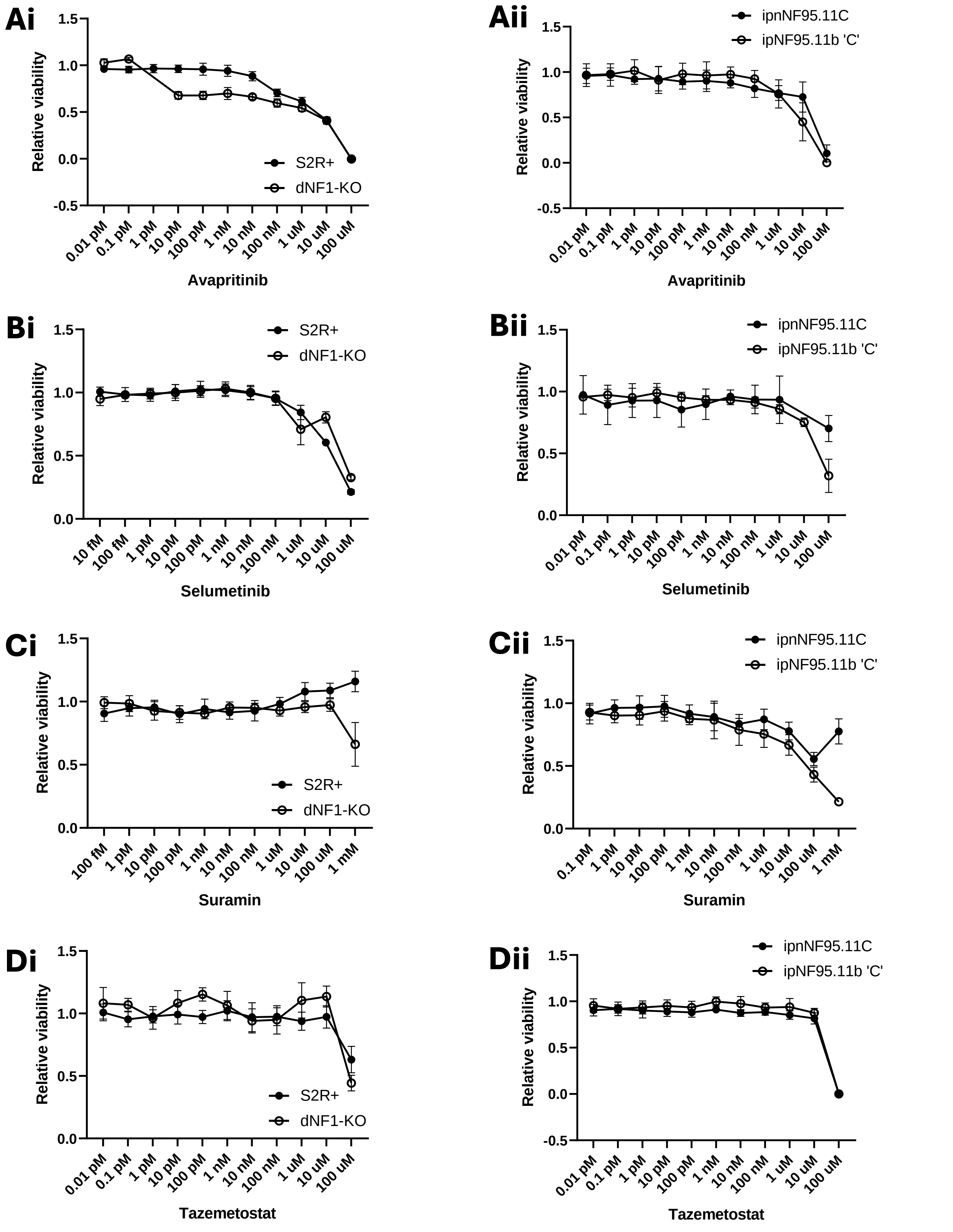

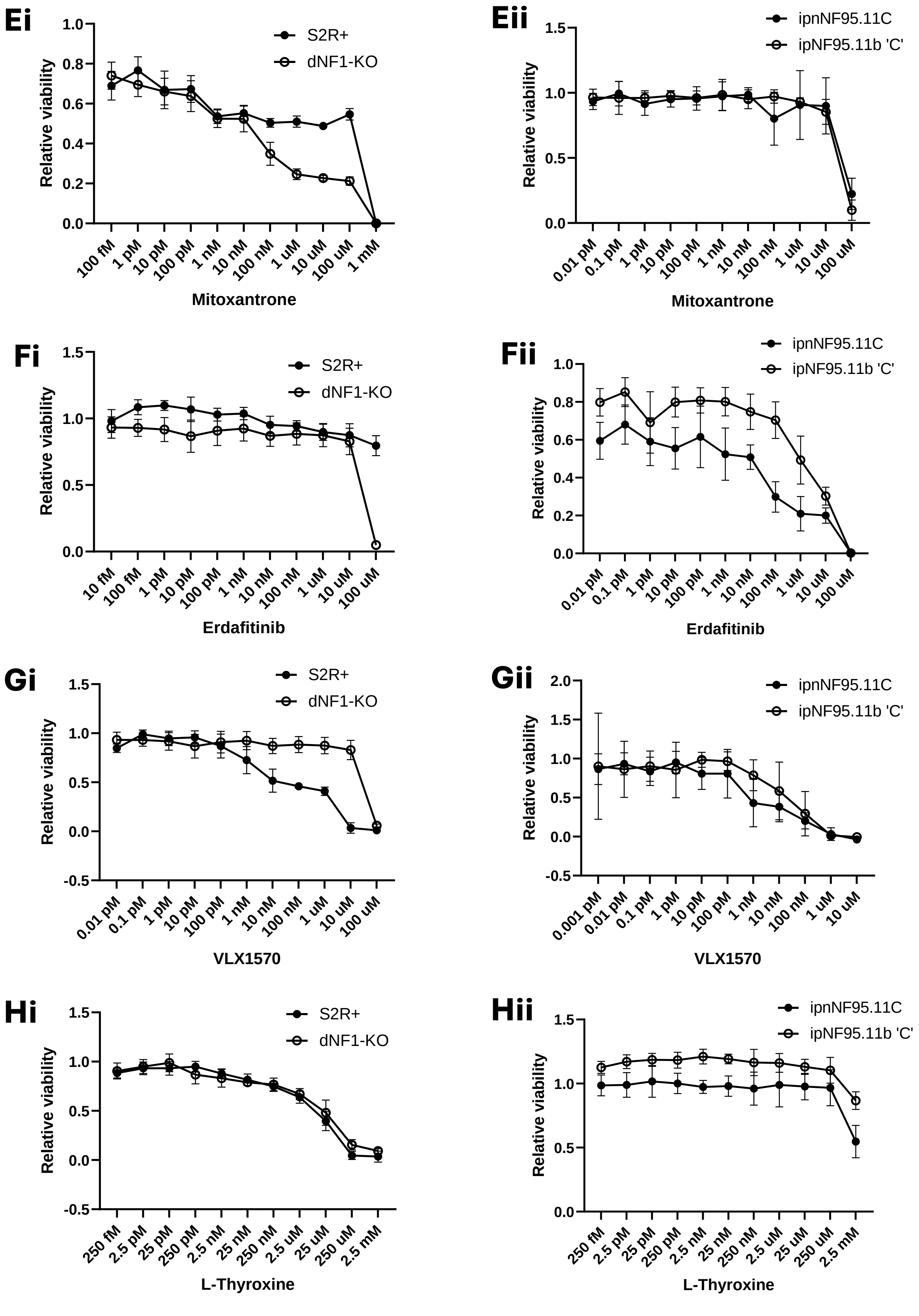

Ei
Relative viability
1.0
0.8
0.6
0.4
0.2
0.0
S2R+
dNF1-KO
100 fM
1 pM
10 pM
100 pM
1 nM
10 nM
100 nM
1 uM
10 uM
100 uM
1 mM
Mitoxantrone
Eii
Relative viability
1.5
1.0
0.5
0.0
ipnNF95.11C
ipNF95.11b 'C'
0.01 pM
0.1 pM
1 pM
10 pM
100 pM
1 nM
10 nM
100 nM
1 uM
10 uM
100 uM
Mitoxantrone
Fi
Relative viability
1.5
1.0
0.5
0.0
S2R+
dNF1-KO
10 fM
100 fM
1 pM
10 pM
100 pM
1 nM
10 nM
100 nM
1 uM
10 uM
100 uM
Erdafitinib
Fii
Relative viability
1.0
0.8
0.6
0.4
0.2
0.0
ipnNF95.11C
ipNF95.11b 'C'
0.01 pM
0.1 pM
1 pM
10 pM
100 pM
1 nM
10 nM
100 nM
1 uM
10 uM
100 uM
Erdafitinib
Gi
Relative viability
1.5
1.0
0.5
0.0
-0.5
S2R+
dNF1-KO
0.01 pM
0.1 pM
1 pM
10 pM
100 pM
1 nM
10 nM
100 nM
1 uM
10 uM
100 uM
VLX1570
Gii
Relative viability
2.0
1.5
1.0
0.5
0.0
-0.5
ipnNF95.11C
ipNF95.11b 'C'
0.001 pM
0.01 pM
0.1 pM
1 pM
10 pM
100 pM
1 nM
10 nM
100 nM
1 uM
10 uM
VLX1570
Hi
Relative viability
1.5
1.0
0.5
0.0
-0.5
S2R+
dNF1-KO
250 fM
2.5 pM
25 pM
250 pM
2.5 nM
25 nM
250 nM
2.5 uM
25 uM
250 uM
2.5 mM
L-Thyroxine
Hii
Relative viability
1.5
1.0
0.5
0.0
ipnNF95.11C
ipNF95.11b 'C'
250 fM
2.5 pM
25 pM
250 pM
2.5 nM
25 nM
250 nM
2.5 uM
25 uM
250 uM
2.5 mM
L-Thyroxine

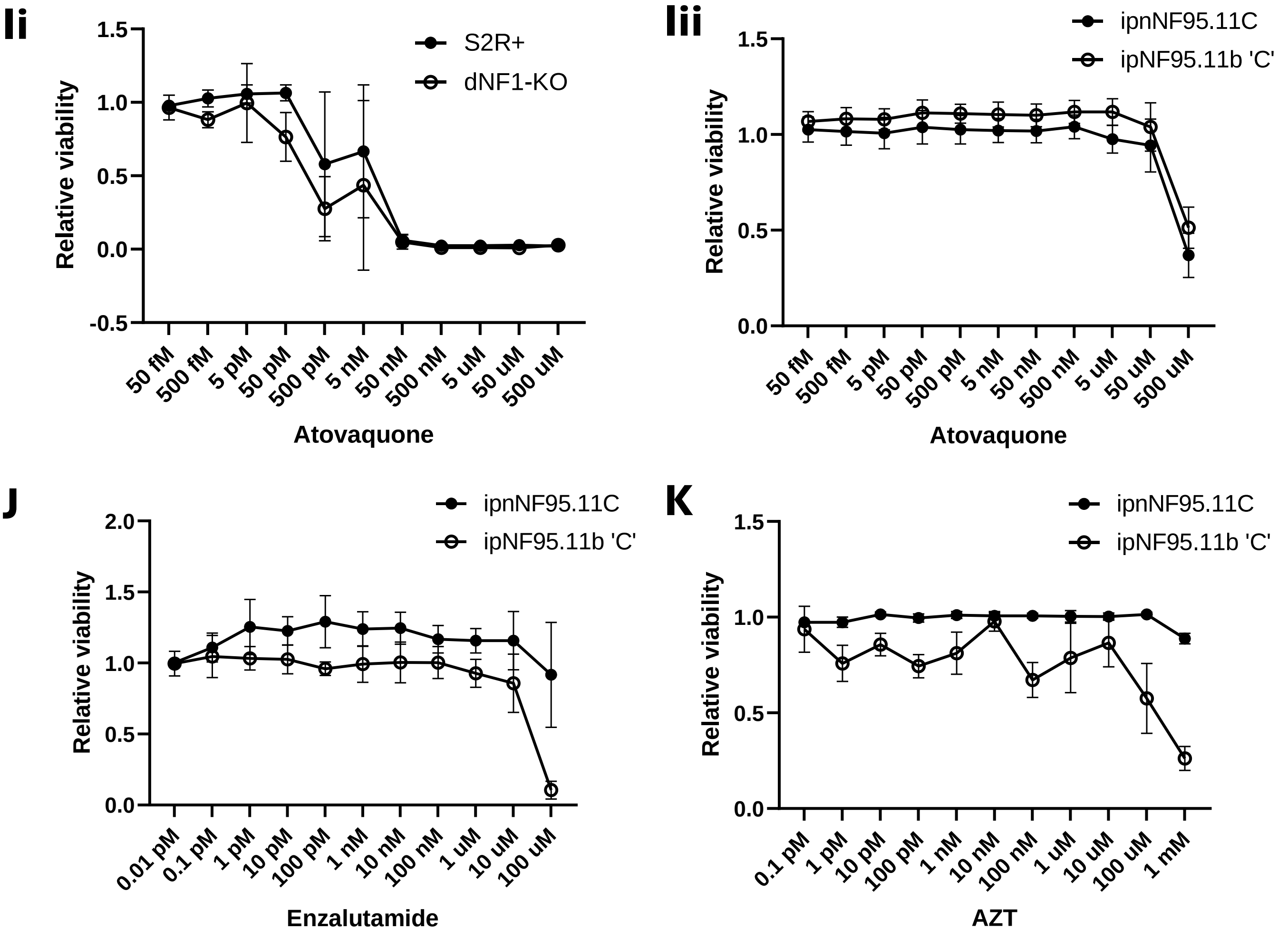
Ii
Relative viability
1.5
1.0
0.5
0.0
-0.5
S2R+
dNF1-KO
50 fM
500 fM
5 pM
50 pM
500 pM
5 nM
50 nM
500 nM
5 uM
50 uM
500 uM
Atovaquone
Iii
Relative viability
1.5
1.0
0.5
0.0
ipnNF95.11C
ipNF95.11b 'C'
50 fM
500 fM
5 pM
50 pM
500 pM
5 nM
50 nM
500 nM
5 uM
50 uM
500 uM
Atovaquone
J
Relative viability
2.0
1.5
1.0
0.5
0.0
ipnNF95.11C
ipNF95.11b 'C'
0.01 pM
0.1 pM
1 pM
10 pM
100 pM
1 nM
10 nM
100 nM
1 uM
10 uM
100 uM
Enzalutamide
K
Relative viability
1.5
1.0
0.5
0.0
ipnNF95.11C
ipNF95.11b 'C'
0.1 pM
1 pM
10 pM
100 pM
1 nM
10 nM
100 nM
1 uM
10 uM
100 uM
1 mM
AZT

**Fig. S2. CellTiter Glo** dose curves for AZT. (A) C8 (Nf1+/-) and C23 (NF1-/-) cells. (B) sNF96.2 NF1-deficient MPNST cells. (C) ST8814 NF1-deficient MPNST cells. *P < 0.05, **P < 0.01.

A

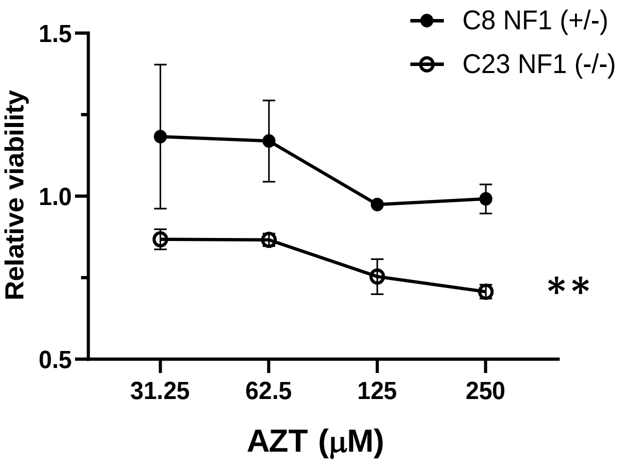


G

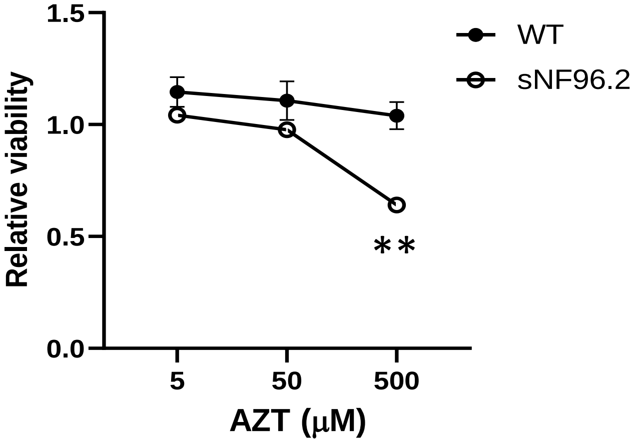


H

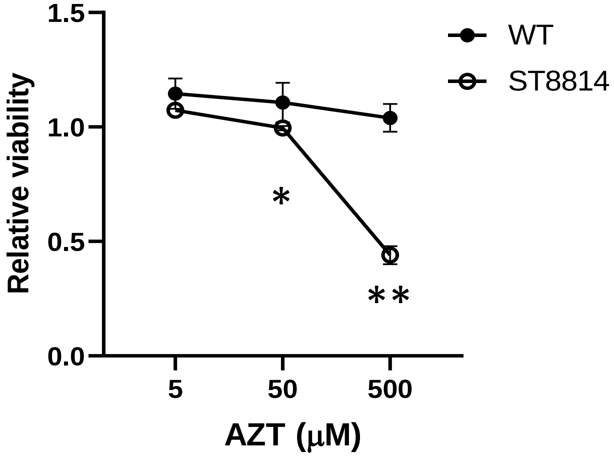

**Fig. S3.** Toxicity of AZT and selumetinib was assessed by weighing mice three times per week. Neither AZT (100 mg/kg) or selumetinib (25 mg/kg) resulted in weight loss in the mice (error bars indicate standard deviation).

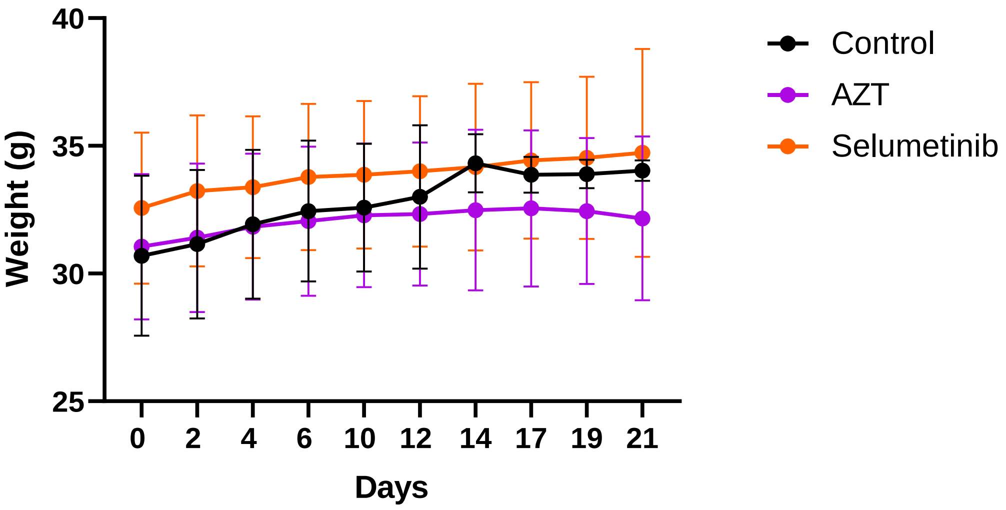

**Fig. S4.** AZT did not show a selective effect on viability in *NF1*-deficient Drosophila S2R+ cells (dNF1-KO), as measured with the CellTiter Glo assay.

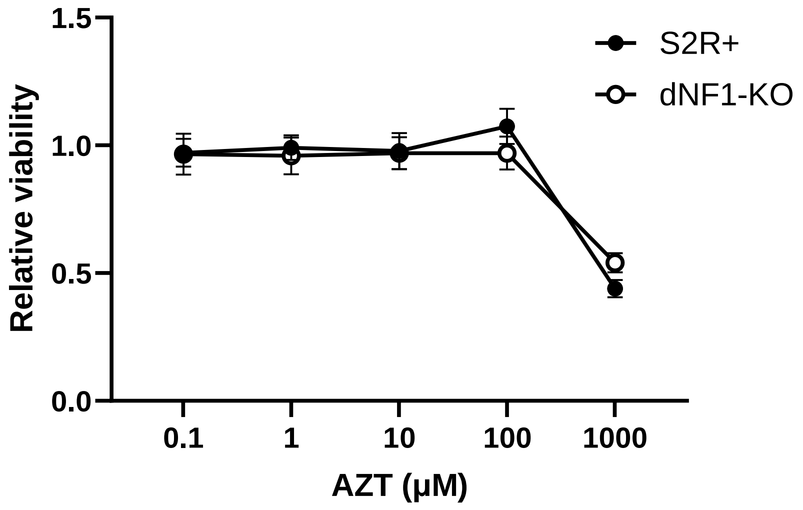

**Fig. S5.** mRNA expression of cell cycle-related genes previously reported to be regulated by AZT in multiple human NF1-deficient cell lines, in addition to a NF1 heterozygous control (ipnNF95.11C). (A) Cyclin-D1; (B) Chk1; (C) Chk2; (D) p14ARF; (E) p53; (F-I) p21. (*P<0.05 vs. control; n=2-5; P values obtained using the Student's t-test; error bars indicate standard deviation).

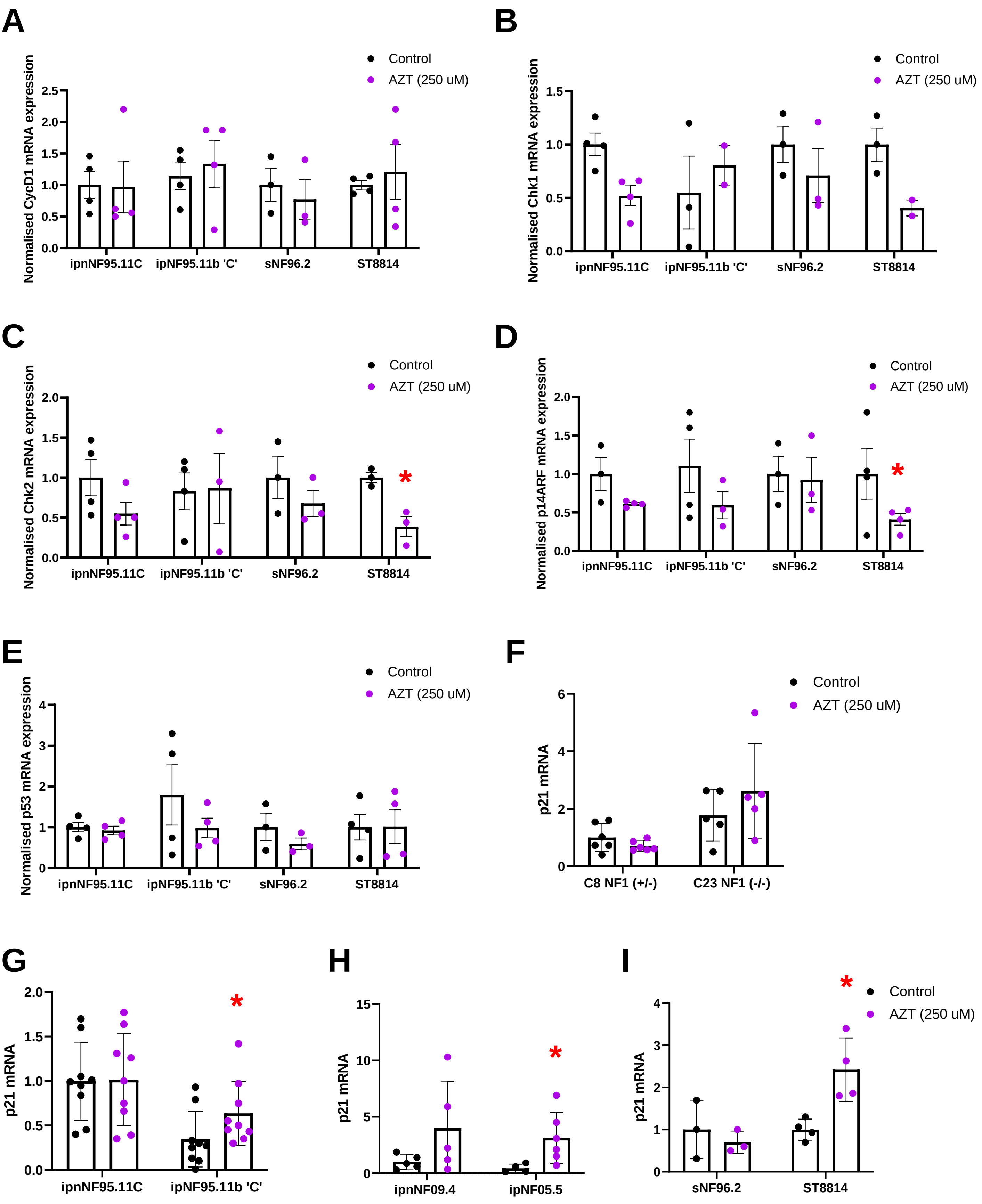
